# Virome of *Culex nigripalpus* at an Alabama aquaculture site reveals diverse insect-specific viruses and the impact of dual bioinformatic pipelines

**DOI:** 10.64898/2025.12.27.696710

**Authors:** Aasma Sharma, Kyle Oswalt, Natalie Wong, Chaoyang Zhao, John F. Beckmann, Kathleen Martin

## Abstract

Mosquitoes are important cosmopolitan insect vectors that threaten humans and host a variety of diseases. Catfish production from Alabama fisheries supports the development of favorable habitats that benefit fish, birds, and insects. Our study found that *Culex nigripalpus* was a major vector associated with catfish ponds. As *Culex nigripalpus* harbors medically important viruses like St. Louis encephalitis, studies often focus on these without exploring the full virus diversity. To address this, we performed an RNA-Seq analysis on *Culex nigripalpus* and compared bioinformatic methodologies, testing two approaches. In the first method (reference-based assembly), the characterized genome of *Culex quinquefasciatus* was used with a mapping cutoff, and assembled contigs were annotated with megablast against the NCBI nucleotide database. In a second method, reads were assembled *de novo*, and the contigs were annotated with a BLAST against the NCBI viral genomic database. In addition, the cross-validation of the contigs generated by the two methods was conducted to identify the common and unique viral contigs, adding support for identification. RNA-Seq analysis identified fifteen different viruses. Both methods identified three common viruses, Merida virus isolate Cx. nigripalpus1 (PQ963471), Hubei mosquito virus 5 isolate Cx. nigripalpus2 (PQ963472), and Zhejiang mosquito virus isolate Cx. nigripalpus3 (PQ963484). The first method also identified ten unique viruses, and the second method generated two unique viruses. Selected viruses were independently validated by RT-PCR amplification of viral genomic regions. Overall, our data suggest that different approaches to bioinformatic analysis complement each other and improve viral genome recovery from metatranscriptomic datasets.

## Introduction

Mosquitoes (Diptera: *Culicidae*) are a cosmopolitan insect group with more than 3600 insect species, many of which are the vectors of parasites, viruses, and bacteria (Nebbak et al., 2021). Mosquitoes have aquatic larval and pupal forms that require water; thus, agricultural fields, wastewater, sewage, bird baths, and anywhere water collects are attractive areas for habitation (Morse et al., 2019). In agricultural settings, this historically encompassed rice paddies and fisheries ecosystems. Mosquitoes from the *Culex* and *Aedes* genera serve as vectors for epidemic diseases from virus groups, including *Flaviviridae*, *Rhabdoviridae*, *Togaviridae* along with viruses in the *order Bunyavirales* (Gómez et al., 2023). Notably members of *Bunyavirales* and some rhabdoviruses are capable of crossing kingdom boundaries, infecting a wide range of hosts. For example, tomato spotted wilt virus (*Orthotospovirus tomatomaculae, Bunyavirales)* infects a wide range of plant hosts and insect hosts such as *Frankliniella occidentealis* (Martin et al., 2025). A less extreme example of a virus that crosses orders is eastern equine encephalitis virus (*Alphavirus eastern*, EEEV), which can infect birds, the mosquito vectors (*Culex spp.)*, humans, and horses (humans and horses are dead-end hosts) (Bingham et. al, 2016; Hughes et. al, 2021). Another virus of concern in the southeastern United States is West Nile virus (*Orthoflavivirus nilense*, WNV), which is vectored by *Culex spp.,* including *Cx. nigripalpus* (Godsey et al., 2013; Richards et al., 2011). These examples highlight the ecological flexibility of such viruses, underscoring their relevance in both agriculture and vector biology.This dynamic becomes especially important in complex ecological regions such as the sub-tropical Gulf Coast of the United States, including Alabama.

Fisheries is a major industry in Alabama, producing 45.1 million catfish in 2023, second only to Mississippi (USDA-NASS). Fish-culture ponds support abundant bird communities, including raptors, wading birds, waterfowl, and songbirds, in the surrounding vegetation (Burr et al., 2020). These systems also harbor diverse reptiles and amphibians such as turtles, snakes, and frogs (Bailey, 1998; Mount, 1975). Many of these vertebrates can act as reservoirs for native, typically non-pathogenic viruses and serve as bloodmeal hosts for a variety of mosquito species. Numerous species of medical importance occur in Alabama, including *Aedes aegypti*, *Ae. albopictus*, *Ae. japonicus*, *Culex pipiens*, *Cx. tarsalis*, *Cx. quinquefasciatus*, and *Anopheles quadrimaculatus*. Additional species have been reported from nearby Tuskegee National Forest, such as *Culiseta melanura*, *Culex restuans*, *Aedes vexans*, *Coquillettidia perturbans*, *Cx. erraticus*, *Cx. peccator*, and other *Culex* and *Ochlerotatus* spp. (Hassan et al., 2003). Overall, more than 60 mosquito species have been documented in Alabama, including *Cx. territans*, *Ochlerotatus triseriatus*, *Cx. salinarius*, *Anopheles punctipennis*, and *An. quadrimaculatus* (Qualls and Mullen, 2006). *Culex nigripalpus* and three *Psorophora* species: *Ps. columbiae*, *Ps. ferox*, and *Ps. ciliata*, have also been recorded from neighboring Macon County (Shaikh et al., 1987). This indicates that *Cx. nigripalpus* is established in the east-central Alabama region that includes our study site in Lee County. The enrichment of species, hosts, and viruses within catfish fisheries is poorly understood, and this study aims to investigate this question.

To investigate novel viruses and microbiomes of mosquitoes in our locale, we focused our study on an ecosystem having three key factors that often contribute to disease outbreaks: proximity to housing developments and human habitation, proximity to undeveloped land such as forested areas, and proximity to aquatic fisheries and aquaculture stations that would amplify mosquito populations. Specifically, in Auburn, a fisheries pond was selected in the northern part of the community, near the Kreher forest and less than a mile from a large housing development. We sought to investigate the mosquito populations in this unique habitat in the context of fisheries agriculture, a valuable cash crop in the state of Alabama. Mosquitoes were collected during early fall 2022 to assess species diversity and to identify the most abundant local vector. From there, our goal was to characterize the virome associated with that dominant species then compare bioinformatic pipeline methodologies. We hypothesized that at least one species of mosquito would predominate aquaculture ponds and that it would carry a multitude of endogenous local viruses. While we did not necessarily expect to detect known human pathogens, given the absence of local outbreak reports, we hypothesized that the prevailing vector would harbor various uncharacterized viruses that we could use to compare the pros and cons of dual bioinformatics RNA-Seq methodologies.

## Materials and Methods

### Collection sites

Mosquitoes utilized for this study were trapped at the Auburn University, E.W. Shell Fisheries center located in Auburn, AL,USA (Figure 1A). We specifically trapped mosquitoes both near the aquaculture pond and near buildings where humans travel close to the pond. The E.W. Shell Fisheries center is a 1600-acre research station consisting of infrastructure and natural habitat. Man-made ponds facilitate studies utilizing diverse species of catfish and trout. The fisheries also provide a habitat for a variety of wildlife including Bald Eagles, Belted Kingfishers, blue herons, wading birds, songbirds, reptiles, and amphibians. The fisheries are located on AL-147, Auburn, Alabama (32.652, -85.486), south of the Kreher Preserve & Nature Center and north of the Auburn campus. This unique location of the fisheries allows it to serve as a novel model representing the interface between a more urban environment to the south and a more natural environment to the north. Its position allows for a more biodiverse ecosystem to be studied for zoonosis.

**Figure 1.**
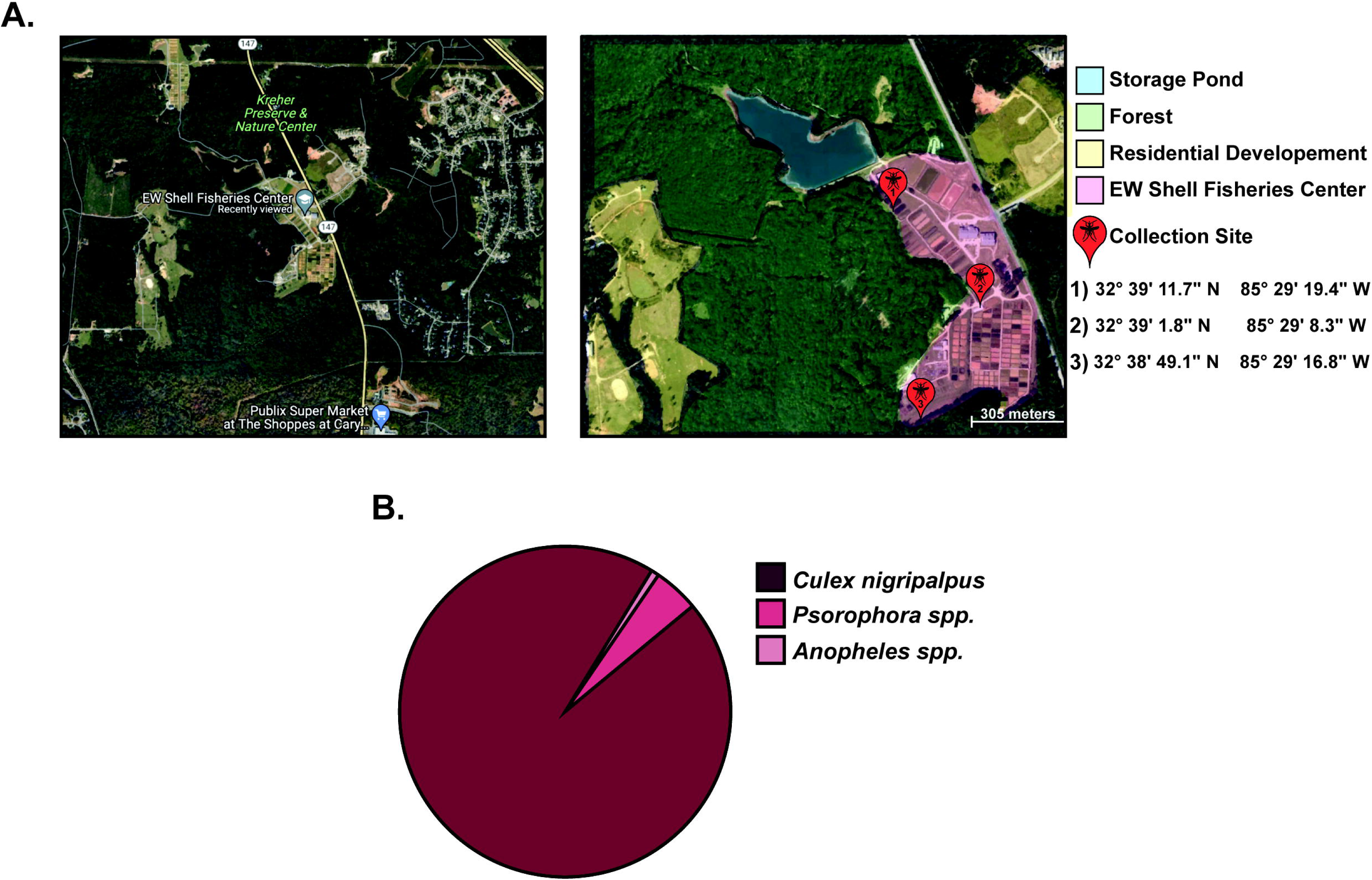
Overview of mosquito sampling of *Culex nigripalpus* collected in E.W. Shell Fisheries between late August to late October in 2022. A) Mapped locations of mosquito collection sites and surrounding areas, including the E.W. Shell Fisheries Center, adjacent forest, storage ponds, and nearby residential developments in Auburn, AL. B) Pie-chart showing the abundance of mosquito species collected during sampling

### Mosquito Collection and Taxonomic Identification

Live adult mosquitoes were collected using Centers for Disease Control (CDC) miniature light traps (Model 512, John W. Hock) baited with CO emitted from approximately 0.9 kg of dry ice per trap. Traps were strategically placed along the edges of the fisheries and forests or at recently drained ponds within E.W. Shell Fisheries (Auburn, AL) to gather a diverse population of mosquitoes. The traps were set overnight from 4 PM until 10 AM the following morning, the collection period extended from late August through the end of October 2022. Upon retrieval, live adult mosquitoes were transported to the laboratory and euthanized by incubation at -20°C for 10 minutes. Subsequently, the mosquitoes were sexed and identified to genus level using a stereomicroscope and a dichotomous key (Darsie and Ward, 2005). Specimens were then grouped by species and sex into pools, with each pool containing a maximum of 20 adult mosquitoes. These pools were submerged in 100% ethanol and stored at -20°C until nucleic acid extraction, as ethanol is a relatively low-cost means of inhibiting RNase activity until extraction, which was performed as soon as the collection interval had passed.

Once the genus level was identified, it was determined that a more thorough evaluation was needed at the genomic level. This was conducted after the initial *de novo* assembly of contigs to identify mosquito sequences. In short: the total contigs from Method Two below were compared to a panel of COI genes: *Culex nigripalpus*: PQ587035.1; *Culex quinquefasciatus*: NC_014574.1; *Culex pipiens pipiens*: NC_015079.1; *Culex erraticus* COI: GenBank KT766417.1. The highest match across the longest region (*Culex nigripalpus*) was selected to make a conclusive identification of the mosquito species collected.

### Viral RNA Extraction

As *Culex nigripalpus* was the predominant species at this location, the study thereafter focused on this species. Female *C. nigripalpus* mosquitoes were removed from ethanol and allowed to dry under a laminar flow hood for 10 minutes. Individuals were then randomly assigned to groups, each comprising 15 to 17 mosquitoes, and placed into 1.5 mL microcentrifuge tubes. This sample size was selected to reduce contaminants for sequencing (Beckmann and Fallon, 2012). Samples were homogenized by pestle in liquid nitrogen, and RNA was extracted from each group with 1 ml of TriZol (Invitrogen, Waltham, MA) according to the manufacturer’s instructions. Modifications to manufacturer’s instructions were implemented to increase the yield and quality of RNA. These steps included repeating the chloroform extraction twice, conducting salt precipitation with 100% isopropanol and 3M sodium acetate (pH 5.2), and two additional cold 70% ethanol washes. Total RNA concentration was quantified by nanodrop (Thermo Scientific, Waltham, MA) and ranged from 1600 ng/µl to 1800 ng/µl. The 260/280 value, which indicates RNA purity, was high, ranging from 2.01 to 2.05, and the 260/230 ratio was 1.88 to 1.92, indicating mild phenol contamination in the sample, as reported by the Nanodrop. Ultimately, RNA samples from four collections with the highest concentrations and purities were pooled to create a single sample for sequencing to capture a snapshot of the viruses present and reduce sequencing costs. Each sample concentration was normalized so that all samples contributed equally to the final pooled sample.

### Ribosomal RNA Depletion and Sequencing

The integrity of total RNA was assessed using ScreenTape according to the manufacturer’s instructions. A subset of the RNA sample was then diluted to a final concentration of 50 ng/µL with molecular-grade water. Subsequently, 1 µL of the diluted RNA sample was mixed with 5 µL of RNA Screen Tape sample buffer. The mixture was briefly vortexed and centrifuged to ensure proper mixing. The Agilent Tape Station instrument (Agilent Technologies) was used to assess RNA quality according to the manufacturer’s instructions. All subsequent steps were conducted according to the manufacturer’s instructions, and the software automatically calculated RNA integrity numbers (RINs) from electropherogram analysis.

Total RNA was sent to SeqCenter (Pittsburgh, Pennsylvania) and prepared for RNA-Seq using their modified ribodepletion pipeline. At the time of RNA extraction and library preparation, the mosquito pool had been identified morphologically as *Culex* spp.; species-level identification as *Culex nigripalpus* was made later from sequence data (see Results). The sample was treated with RNase-free DNase (Invitrogen, Waltham, MA), and ribodepletion was carried out using custom probes designed and applied by SeqCenter. Probes were designed to target conserved regions of *Culex* spp. ribosomal RNA (5.8S, 28S, and 18S rRNA). Reference 18S and 28S sequences from *Culex* spp., including representatives of the Melanoconion subgenus (Burkett-Cadena et al., 2022; Koh et al., 2023), were aligned and used for probe design to maximize depletion efficiency across *Culex* mosquitoes. The library was then prepared using an Illumina Stranded Total RNA Prep Ligation kit with 10-bp unique dual indices (UDI) (Illumina, San Diego, CA). Sequencing was performed on a NovaSeq X Plus (Illumina, San Diego, CA), generating ∼50 million paired-end 150-bp reads. Demultiplexing, quality control, and adapter trimming were performed using bcl-convert (v4.1.5) by SeqCenter.

### Bioinformatic analysis pipelines and Cross-Validation

Once the sequences were received, the two different analysis pipelines were followed (Figure 2). The methods are briefly described below. In Method One, total reads were first mapped first to a mosquito reference genome. Unmapped reads were more likely to contain viruses, therefore, they were subsetted and assembled using *de novo* assembly and the contigs were utilized to BLAST against the universal NCBI database. In Method Two, total reads were utilized for *de novo* assembly and the contigs were blasted against the virus NCBI viral genomic database (viral.1.1.genomic.fna) (NCBI Virus, n.d.) (Figure 2).

**Figure 2.**
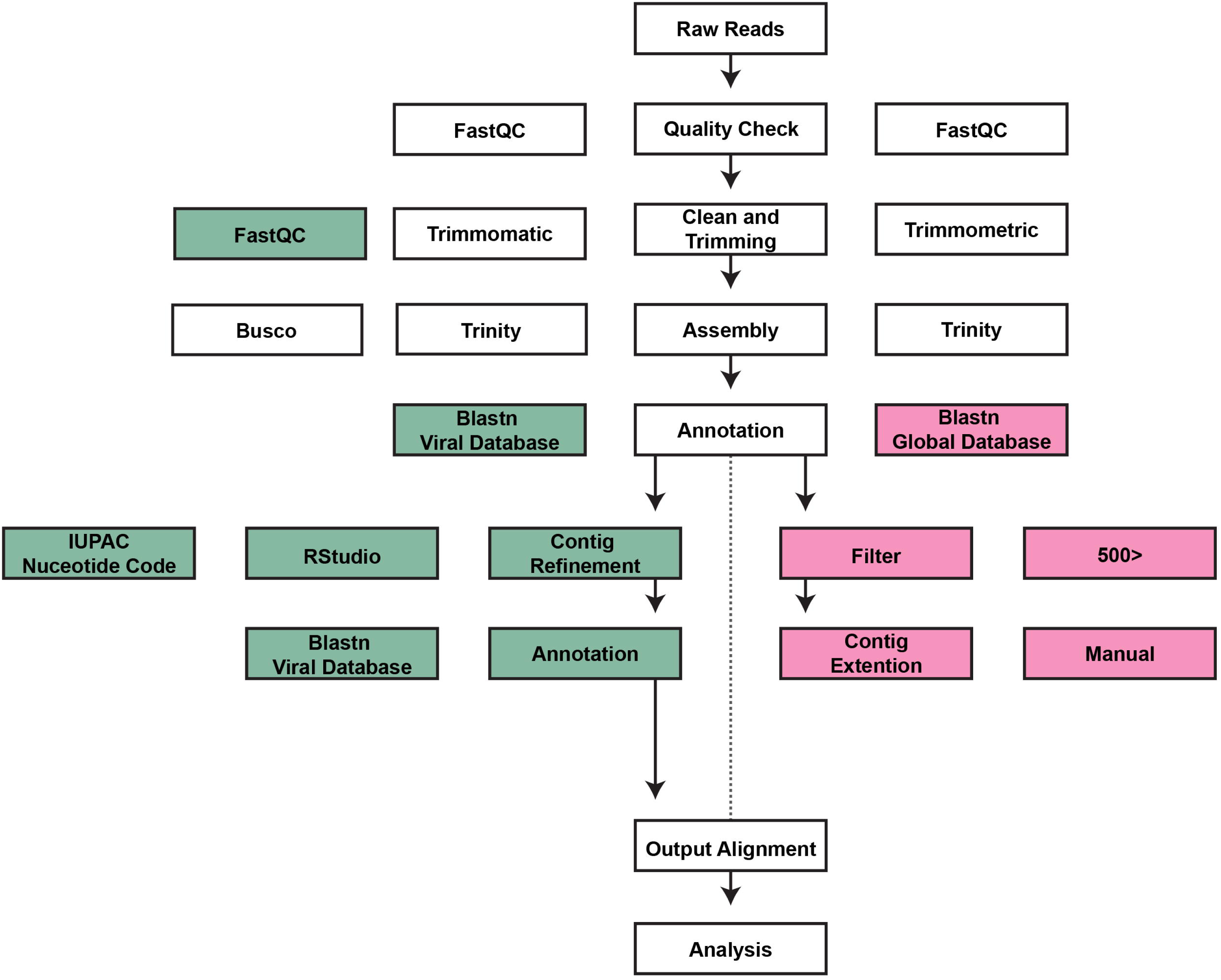
Bioinformatic pipeline: Dual bioinformatic pipeline for the virome analysis of *Culex nigripalpus* sampled in E.W. Shell Fisheries in Auburn, AL in 2022

In detail, for Method One, sequence quality was assessed using FastQC (version 0.74), and reads were trimmed using Trimmomatic (Galaxy version 0.39+galaxy2) with default settings to remove low-quality bases and reads. Then the reads were mapped using Bowtie2 (Langmead et al., 2009a), (Langmead et al., 2009b) against the complete reference genome of *Culex quinquefasciatus* (GCF_015732765.1) using the default parameters on the galaxy bioinformatics platform (usegalaxy.org) (Community, 2024). The reads that aligned to the reference genome were filtered out. Those that did not align to the reference genome were subsetted and assembled using using the default settings on Trinity Version 2.15.1 (Grabherr et al, 2014), on the same platform (Community, 2024). Assembled contigs were separated based on the size of contigs using Filter sequences by length (Galaxy Version 1.2) into 0-400 bp, 400-1000bp and >1000 bp. These contigs were blasted using NCBI BLAST+ BLASTN (Cock et al., 2015) (Camacho et al., 2009a) against NCBI-NT 2023 universal database. Following final assembly, viral contigs <400 bp were excluded. For each virus, the longest contig was used as the representative sequence, with nucleotide identity (%) used to break ties among contigs of similar length (Batson et al., 2021). When needed, the longest contig was utilized as the representative scaffold and extended only if additional contigs with the same viral annotation showed consistent overlap. For unique contigs, the longest contig with high amino acid identity (%) annotation was considered to ensure the sensitivity of the virus during detection and reduce the need for manual genome extension (Deng et al., 2015; Deng and Delwart, 2021).

In detail, for Method Two, raw sequencing data processing followed the same methods as Method One without reference alignment and subsetting. Subsequently, high-quality reads were then assembled *de novo* using Trinity (Galaxy version 2.11.0) (Grabherr et al., 2011) with default parameters. The resulting contigs were queried against the NCBI viral genomic database (viral.1.1.genomic.fna) using BLASTN (BLAST+ 2.16.0) with an E-value threshold of 1e-5. Contigs with significant viral matches were clustered by shared annotation. For each viral hit, the longest contig with a high identity was retained as a representative sequence. Where overlapping hits were present, consensus sequences were generated using IUPAC nucleotide codes to reflect polymorphic base positions across contigs.

To determine the similarity of the sequences from each method to each other, command-line NCBI+ tools were utilized. Specifically, local BLAST commands were used to cross-compare contigs predicted by each pipeline (Camacho et al., 2009b). To determine transcript quantification for the full-genome top hits in this dataset, bias-aware transcript quantification was performed using Kallisto (kallisto_linux-v0.48.0) (Bray et al., 2016).

### Validation of the Sequences

To determine whether the contigs generated in the bioinformatic analysis were consistent with those found *in vivo*, primers were designed for a series of candidate viruses using NCBI’s Primer-Blast tool. Contigs from Method One were used to design PCR primers to validate as many detected viruses as possible. When multiple contigs were annotated as the same virus, primers were sometimes designed from different contigs to help confirm that the assembly was correct. In short, each contig was analyzed using this tool, with default settings except for minimum and maximum contig lengths of 500 and 2000 bases, respectively. The top candidate primer pair was selected for each contig. When possible, multiple pairs were designed for each virus based on different contigs. As the concentration was over 1.5 μg/mL, the total RNA was diluted from 2 μL to 10 μL to facilitate pipetting. The cDNA was synthesized using two different kits, the RevertAid kit (Thermo Fisher Scientific) and the Verso cDNA kit (Thermo Fisher Scientific), following the instructions provided by the companies, using a random hexamer: OligodT primer mix at a 3:1 ratio as described in the Verso cDNA kit and one μL of the diluted total RNA. For both enzymes, the same reverse primer mix and the RNA were incubated at 65 °C for ten minutes before adding the additional components as described in the kit directions. They were then incubated at 42 °C for one hour. The RevertAid was deactivated by incubation at 70 °C for five minutes, and the Verso was incubated at 95 °C for two minutes.

The cDNA was used for PCR with Phusion polymerase (New England Biolabs) and specific primers provided in Supplementary Table I, following the kit instructions. As all primer pairs had similar TMs (ranging from 57-63 °C), the lowest was selected at 57 °C. The extension time was set to 1 minute and 30 seconds to accommodate larger amplifications for longer contigs. The amplification followed the other default settings for Phusion as provided by the kit instructions. Once complete, the PCR products were analyzed by running 5 μL of the 50 μL reaction mix on a 1% Agarose gel using SyberSafe (Invitrogen, Inc) for 45min at 80V.

### Phylogenetic Analysis

To understand the phylogenetic relationship of Merida virus isolate Cx.nigripalpus1 (PQ963471) to other complete or nearly complete Merida virus isolate available in the NCBI database, a maximum likelihood phylogenetic tree was generated using MEGA 12.1 (Kumar et al., 2024; Stecher et al., 2025) and the tree was visualized using ITOL tree maker (Letunic & Bork, 2024). The accession number of the virus >10,000kb were used to generate the trees: NC_040599.1, MW434769.1, MW434773.1, OM817546.1, MW434771.1, MW434770.1, MW434772.1, MH188000.1, MH310083.1, OL700074.1, MT577803.1, NC040532.1, MF882997.1, OQ725971.1, OQ725976.1, OQ725974.1, OQ725980.1, OQ725972.1, PQ773030.1, MK440621.1, PQ773029.1, OQ725979.1, OQ725979.1, OQ725977.1, OQ725977.1, OQ725978.1, OQ725978.1, and OQ725973.1.

### Analysis of Unannotated Contigs

As a proportion of contigs were not annotated in the BLASTN approaches, the non-annotated contigs were separated for each of the two methods into separate FASTA files using Python 3.13.5. Once separated, each file, M1.unannotated and M2.unannotated, was used for taxonomic analysis to identify any further viral contigs. Taxonomic analysis of unclassified contigs was conducted following the DIAMOND-MEGAN protocol previously described (Zhao et al., 2025). Contig sequences were initially searched against the NCBI non-redundant database, downloaded on August 28, 2023, using DIAMOND (v2.1.8) in BLASTX mode. The output was then processed for taxonomic assignment using the MEGAN database (megan-map-Feb2022.db) in long-read mode via MEGANIZER, a tool included in the MEGAN package (v6.25.3). Finally, sequences classified under the taxonomic label “Virus” were retrieved in MEGAN’s interactive mode.

To determine whether avian haemosporidian parasites were present in *Culex nigripalpus* from our site, we specifically screened for *Plasmodium* spp., including *Plasmodium hermani*, which has previously been associated with *Cx. nigripalpus* (Forrester et al., 1980). As sequence data for *P. hermani* are not available, we compiled a reference panel of avian *Plasmodium* sequences from GenBank (KC142195 *Plasmodium juxtanucleare* isolate GALLUS08; KP326567 *P. juxtanucleare*; KT290910 *P. juxtanucleare* isolate PjMH04; KY653773 *P. relictum* clone PCc235-13C; KY653774 *P. relictum* clone Peng14-121Br2AF; KY653801 *P. elongatum* clone PCc370-14C1; NC_012426 *P. relictum*). Assembled mosquito contigs were queried against this reference panel using BLASTN using default parameters, and hits were examined for sequence similarity to avian *Plasmodium*; this approach was intended to detect contigs showing homology to *Plasmodium* spp., even if not an exact match to any single reference sequence.

## Results

### *Culex nigripalpus* is Amplified in Alabama Aquaculture

Between August and October 2022, a total of 11 mosquito collections were assembled in multiple locations within E.W. Shell Fisheries in Auburn, Alabama. Mosquitoes were collected from ecotones between forested areas and fisheries, as well as recently drained ponds, to enhance biodiversity surveillance. From these collections, a total of 237 female mosquitoes were collected, sexed, and morphologically identified to genus and species with the use of a stereomicroscope and taxonomic key (Table 1). The results revealed the presence of 3 unique genera and species at the fisheries, *Culex spp.*, *Psorophora* spp., and *Anopheles crucians*. Once the genetic sequences of the samples were evaluated, it was determined to be *Culex nigripalpus* Theobald, with 97-100% of the mitochondrial genome covered by contigs from our sample (Table 2). *Cx. nigripalpus* was the most abundant of the three mosquito species collected, comprising 97% (230) of the total specimens, followed by *Psorophora spp*. (2.5%, 6) and *Anopheles crucians* (0.4%, 1) (Table 1). The presence of these species was not unexpected given their widespread distribution in the southeastern region of North America (Darsie & Ward, 2016). Still, the data indicate that *Culex nigripalpus* breeds in and is amplified by aquaculture, potentially due to the large numbers of birds nearby and the semi-urban environment, which provides a variety of other hosts as well. Due to this, *Culex nigripalpus* was selected for subsequent virome analysis (Table 1, 2, Figure 1B).

**Table 1.** Mosquito Collection at the Fisheries. This Table depicts the mosquito collections taken from E.W. Shell Fisheries between late August to Late October in 2022.

| Sampling Date | Species of Mosquito collected | Male/Female | Number |
| --- | --- | --- | --- |
| 8/23/22 | *Culex nigripalpus* | Female | 21 |
| 8/31/22 | *Culex nigripalpus* | Female | 12 |
| 9/1/22 | *Anopheles Crucians* | Female | 1 |
| 9/7/22 | *Culex nigripalpus* | Female | 8 |
| 9/7/22 | *Culex nigripalpus* | Female | 5 |
| 9/7/22 | *Psorophora spp.* | Female | 5 |
| 9/21/22 | *Culex nigripalpus* | Female | 80 |
| 9/21/22 | *Culex nigripalpus* | Female | 4 |
| 10/12/22 | *Culex nigripalpus* | Female | 20 |
| 10/26/22 | *Culex nigripalpus* | Female | 80 |
| 10/26/22 | *Psorophora spp.* | Female | 1 |
| Total |  |  | 237 |

**Table 2.**
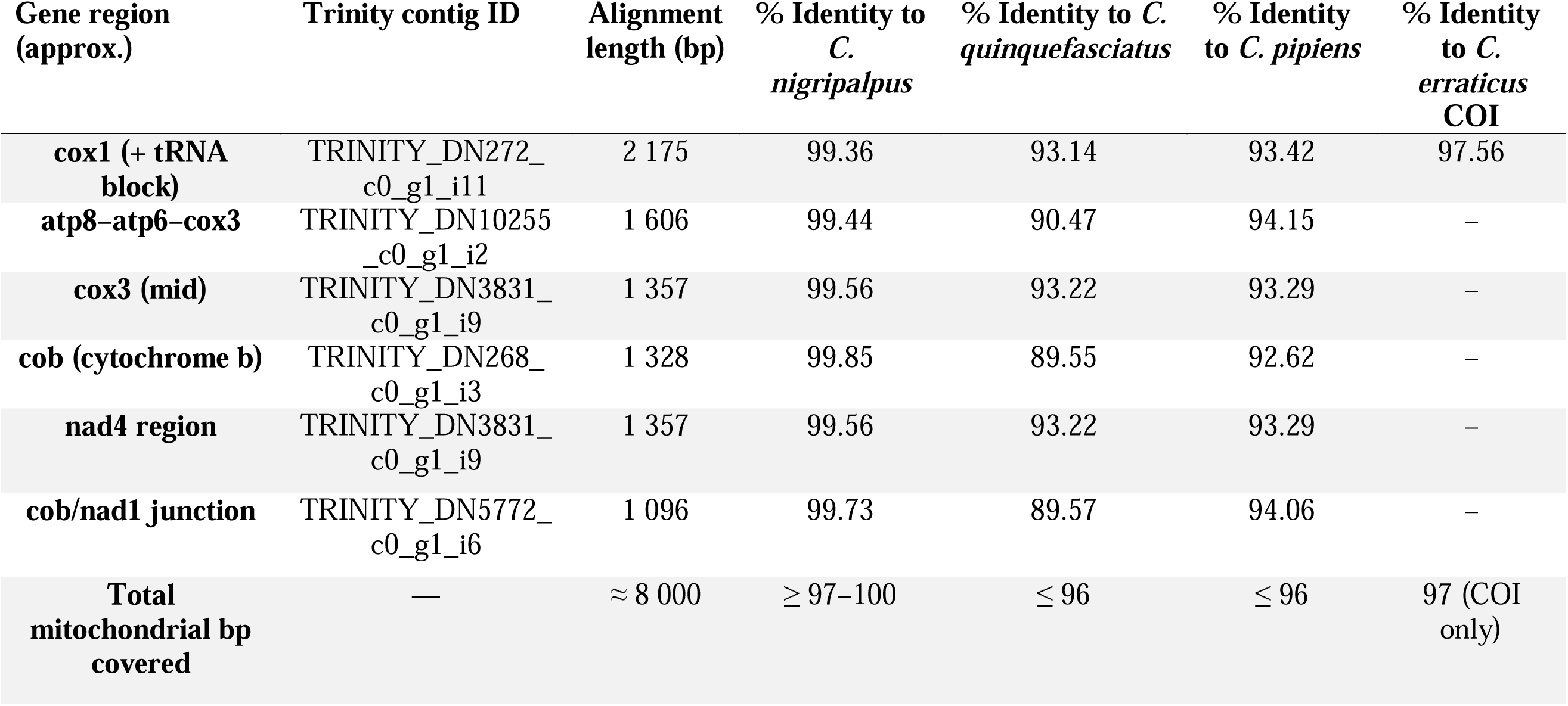
Top BLASTN alignments of host mitochondrial contigs against Culex reference mitogenomes. (*Culex nigripalpus*: PQ587035.1; *Culex quinquefasciatus*: NC_014574.1; *Culex pipiens pipiens*: NC_015079.1; *Culex erraticus* COI: GenBank KT766417.1).

| Gene region<br>(approx.) | Trinity contig ID | Alignment<br>length (bp) | % Identity to<br>*C.*<br>*nigripalpus* | % Identity to *C.*<br>*quinquefasciatus* | % Identity<br>to *C. pipiens* | % Identity<br>to *C.*<br>*erraticus*<br>COI |
| --- | --- | --- | --- | --- | --- | --- |
| **cox1 (+ tRNA**<br>**block)** | TRINITY_DN272_<br>c0_g1_i11 | 2 175 | 99.36 | 93.14 | 93.42 | 97.56 |
| **atp8–atp6–cox3** | TRINITY_DN10255<br>_c0_g1_i2 | 1 606 | 99.44 | 90.47 | 94.15 | – |
| **cox3 (mid)** | TRINITY_DN3831_<br>c0_g1_i9 | 1 357 | 99.56 | 93.22 | 93.29 | – |
| **cob (cytochrome b)** | TRINITY_DN268_<br>c0_g1_i3 | 1 328 | 99.85 | 89.55 | 92.62 | – |
| **nad4 region** | TRINITY_DN3831_<br>c0_g1_i9 | 1 357 | 99.56 | 93.22 | 93.29 | – |
| **cob/nad1 junction** | TRINITY_DN5772_<br>c0_g1_i6 | 1 096 | 99.73 | 89.57 | 94.06 | – |
| **Total**<br>**mitochondrial bp**<br>**covered** | — | ≈ 8 000 | ≥ 97–100 | ≤ 96 | ≤ 96 | 97 (COI<br>only) |

### RNA-Seq Processing Statistics: Reference-Based Assembly (Method One)

The effectiveness of the reference-based approach in identifying non-host viral sequences was evaluated by examining the characteristics of the raw reads and their alignment results to the closest available reference genome. The basic strategy is to use alignment against a reference genome to assemble and subset contigs. Contigs not aligning with the host mosquito genome are then further analyzed for viral content. A total of 158,709,252 reads (79,354,626 paired-end forward and reverse reads each) comprised 10.5 Gbp of bases. Sequence lengths ranged from 35 bp to 150 bp, with a GC content of 45% and an average Phred score of 33. The BioProject ID for the raw read sequences is PRJNA1227547 (submission ID: SAMN46988613). Due to the absence of a *Culex nigripalpus* genome in the NCBI database, mapping was conducted against the *Culex quinquefasciatus* genome. A total of 16.72% of the sequences aligned to the reference genome (GCF_015732765.1), while the remaining 83.28% did not. This is not unexpected, because our libraries contain RNA from the mosquito host as well as viruses, symbionts, microbiota, and environmental contaminants. Additionally, the reference genome is from a different species (*Cx. quinquefasciatus*) than our focal mosquito, *Cx. nigripalpus* (both in the subgenus *Culex*). These data indicate that reference-based approaches work best when a published genome of the target species is available; using “close” relative genomes is not the most robust approach because alignment is generally low causing host reads to contaminate the non-aligned read subset; furthermore, sequence divergence can occur at rates depending on evolutionary divergence and exacerbate the contamination (Payá-Milans et al., 2018). As such, 16.7% of reads (mosquito-derived) were excluded from further analysis, and the remaining unaligned reads were used for downstream assembly. To minimize read redundancy, Trinity was used for transcriptome assembly, which yielded 1,006,209 non-mapped transcripts with a GC content of 41.27%, an average contig length of 580.25 bp, and a total of 583,848,302 assembled bases. Among these, 552,341 contigs were <400 bp, 325,249 contigs ranged from 400–1000 bp, and 128,619 contigs were >1000 bp (Figure 3A). Looking more closely, 222,064 contigs had matches with Megablast compared to the majority that did not. Of those that matched Genbank, only 857 contigs mapped to viral keywords such as “virus, viral, phage or viridae”, while the majority of contigs matched to non-viral keywords such as “culex (176,736), anopheles (2,743), aedes (2,414) and mosquito (21)” suggesting a large portion of the contigs were of insect origin and were not accurately removed from the set, due to genomic differences against the reference genome (Figure 3B). After final assembly and filtering, a refined set of 12 distinct viruses was characterized (Table 3).

**Figure 3.**
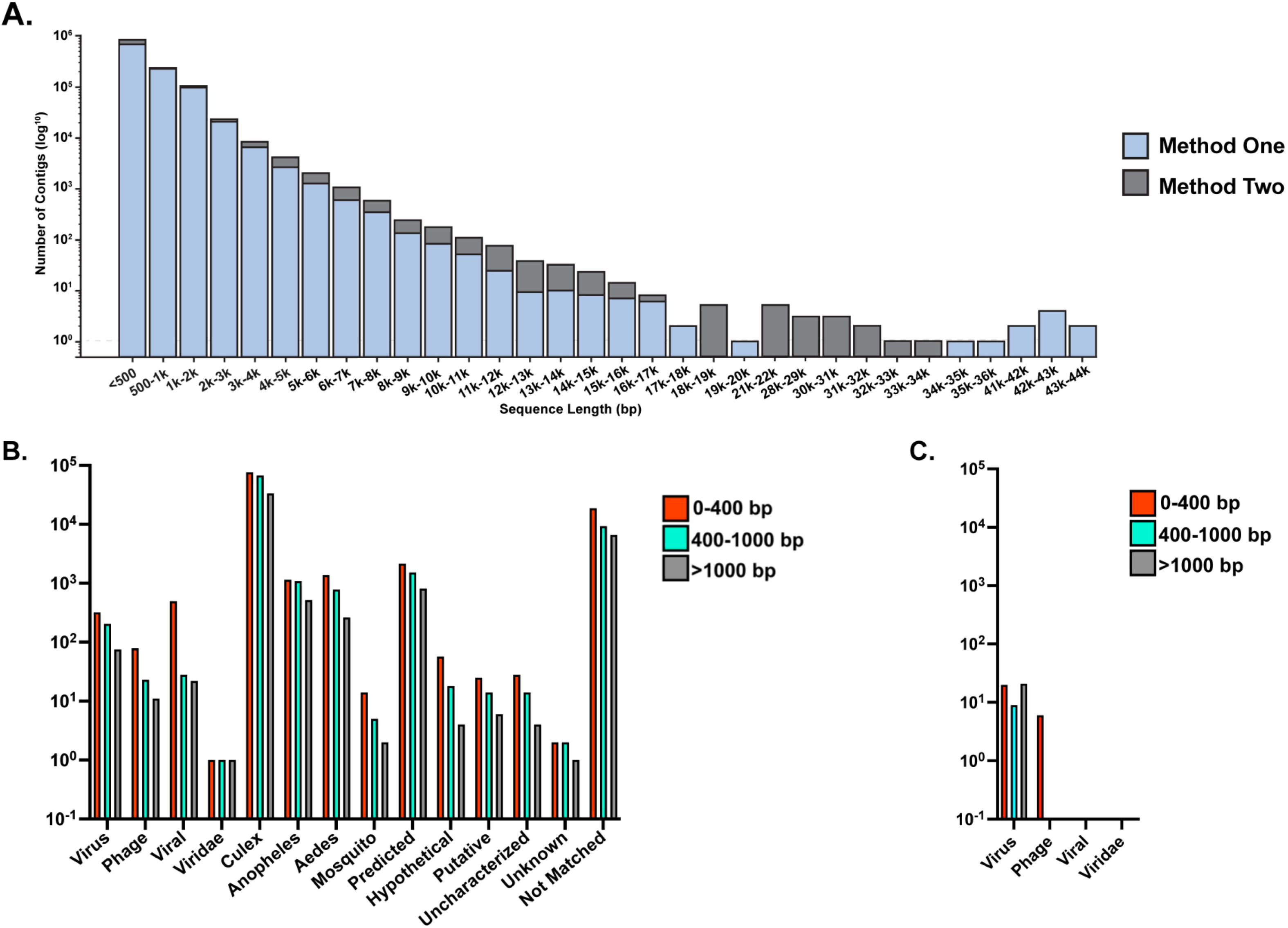
Contig statistics: A) Statistics of contig size after assembly by Method One and Method Two after assembly by Trinity Version 2.15.1 (Grabherr et al, 2014) and Trinity (Galaxy version 2.11.0) (Grabherr et al., 2011) respectively; Blue bar represents the contig length from Method One and Grey bar represents the contig length from Method Two B)Contig annotation after blasting against Universal Megablast by NCBI BLAST+ BLASTN (Cock et al., 2015) (Camacho et al., 2009a) against NCBI-NT 2023 universal database in Method One C) Contig annotation after blasting against NCBI viral genomic database (viral.1.1.genomic.fna) using BLASTN (BLAST+ 2.16.0); Red bar represents the contigs of size 0-400 bp, Blue bar represents the contigs of size 400-1000bp and grey bar represents the contigs of size >1000 bp; In figure B and C, Y axis represents the number of contigs

**Table 3.**
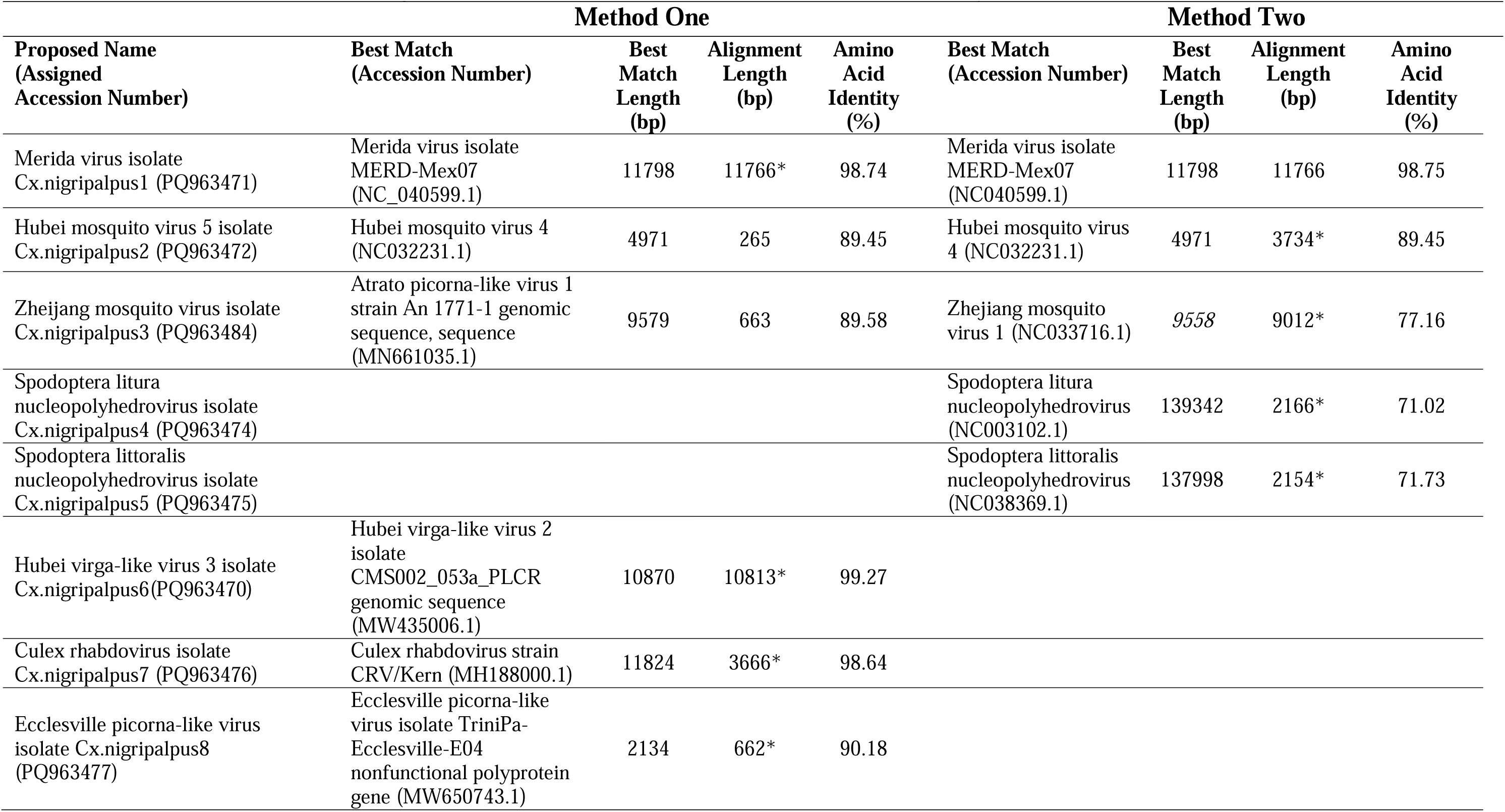
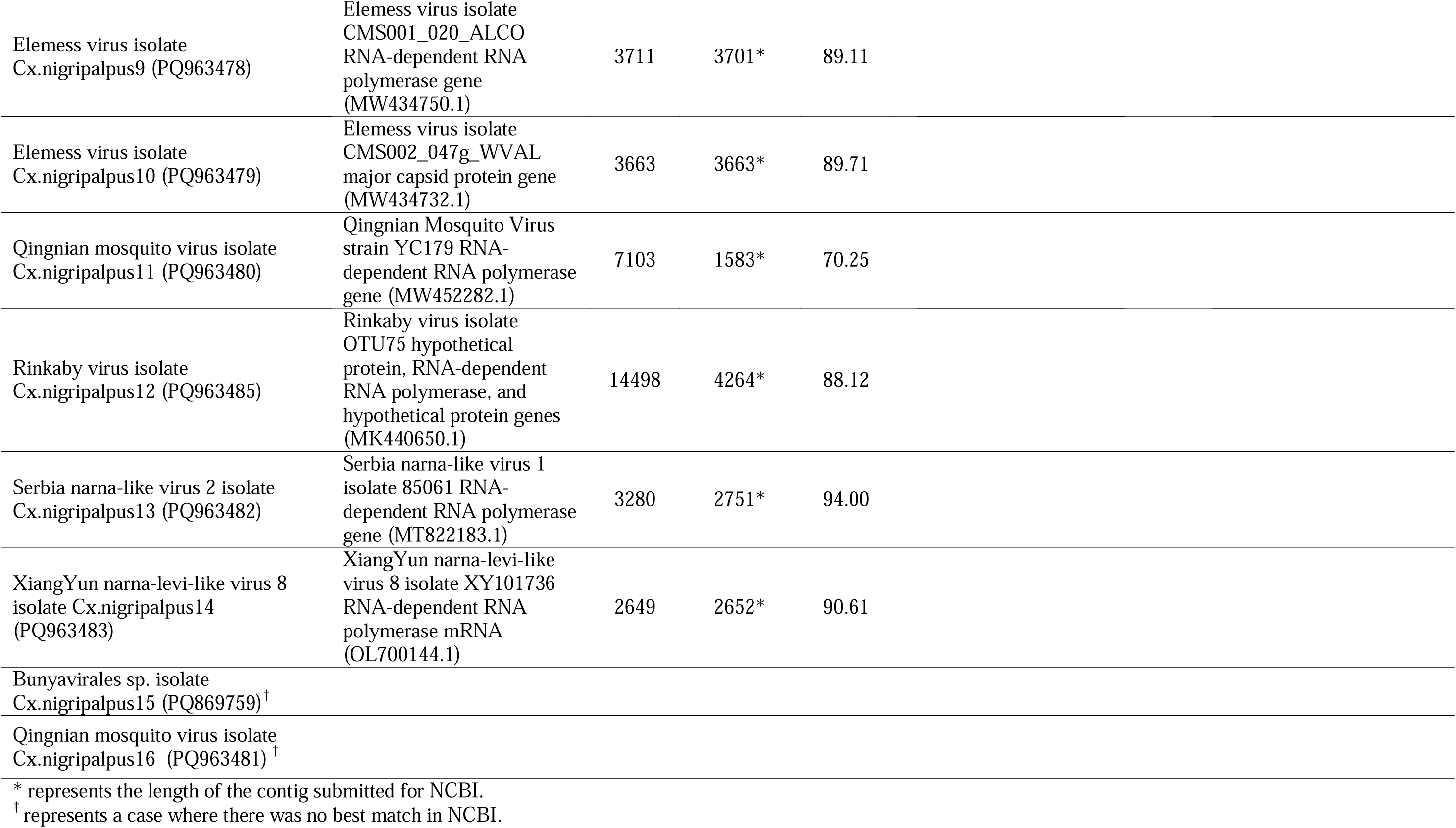
Diversity of viruses. The table shows the diversity of viruses from RNA-Seq analysis of *Culex nigripalpus* collected in E.W. Shell Fisheries between late August to Late October in 2022; RNA: Ribo-Nucleic Acid by dual bio-informatic methods

| Method One |  |  |  |  | Method Two |  |  |  |
| --- | --- | --- | --- | --- | --- | --- | --- | --- |
| Proposed Name<br>(Assigned<br>Accession Number) | Best Match<br>(Accession Number) | Best<br>Match<br>Length<br>(bp) | Alignment<br>Length<br>(bp) | Amino<br>Acid<br>Identity<br>(%) | Best Match<br>(Accession Number) | Best<br>Match<br>Length<br>(bp) | Alignment<br>Length<br>(bp) | Amino<br>Acid<br>Identity<br>(%) |
| Merida virus isolate<br>Cx.nigripalpus1 (PQ963471) | Merida virus isolate<br>MERD-Mex07<br>(NC_040599.1) | 11798 | 11766* | 98.74 | Merida virus isolate<br>MERD-Mex07<br>(NC040599.1) | 11798 | 11766 | 98.75 |
| Hubei mosquito virus 5 isolate<br>Cx.nigripalpus2 (PQ963472) | Hubei mosquito virus 4<br>(NC032231.1) | 4971 | 265 | 89.45 | Hubei mosquito virus<br>4 (NC032231.1) | 4971 | 3734* | 89.45 |
| Zhejiang mosquito virus isolate<br>Cx.nigripalpus3 (PQ963484) | Atrato picorna-like virus 1<br>strain An 1771-1 genomic<br>sequence, sequence<br>(MN661035.1) | 9579 | 663 | 89.58 | Zhejiang mosquito<br>virus 1 (NC033716.1) | 9558 | 9012* | 77.16 |
| Spodoptera litura<br>nucleopolyhedrovirus isolate<br>Cx.nigripalpus4 (PQ963474) |  |  |  |  | Spodoptera litura<br>nucleopolyhedrovirus<br>(NC003102.1) | 139342 | 2166* | 71.02 |
| Spodoptera littoralis<br>nucleopolyhedrovirus isolate<br>Cx.nigripalpus5 (PQ963475) |  |  |  |  | Spodoptera littoralis<br>nucleopolyhedrovirus<br>(NC038369.1) | 137998 | 2154* | 71.73 |
| Hubei virga-like virus 3 isolate<br>Cx.nigripalpus6(PQ963470) | Hubei virga-like virus 2<br>isolate<br>CMS002_053a_PLCR<br>genomic sequence<br>(MW435006.1) | 10870 | 10813* | 99.27 |  |  |  |  |
| Culex rhabdovirus isolate<br>Cx.nigripalpus7 (PQ963476) | Culex rhabdovirus strain<br>CRV/Kern (MH188000.1) | 11824 | 3666* | 98.64 |  |  |  |  |
| Ecclesville picorna-like virus<br>isolate Cx.nigripalpus8<br>(PQ963477) | Ecclesville picorna-like<br>virus isolate TriniPa-Ecclesville-E04<br>nonfunctional polyprotein<br>gene (MW650743.1) | 2134 | 662* | 90.18 |  |  |  |  |

|  |  |  |  |  |
| --- | --- | --- | --- | --- |
| Elemess virus isolate<br>Cx.nigripalpus9 (PQ963478) | Elemess virus isolate<br>CMS001_020_ALCO<br>RNA-dependent RNA<br>polymerase gene<br>(MW434750.1) | 3711 | 3701* | 89.11 |
| Elemess virus isolate<br>Cx.nigripalpus10 (PQ963479) | Elemess virus isolate<br>CMS002_047g_WVAL<br>major capsid protein gene<br>(MW434732.1) | 3663 | 3663* | 89.71 |
| Qingnian mosquito virus isolate<br>Cx.nigripalpus11 (PQ963480) | Qingnian Mosquito Virus<br>strain YC179 RNA-dependent RNA polymerase<br>gene (MW452282.1) | 7103 | 1583* | 70.25 |
| Rinkaby virus isolate<br>Cx.nigripalpus12 (PQ963485) | Rinkaby virus isolate<br>OTU75 hypothetical<br>protein, RNA-dependent<br>RNA polymerase, and<br>hypothetical protein genes<br>(MK440650.1) | 14498 | 4264* | 88.12 |
| Serbia narna-like virus 2 isolate<br>Cx.nigripalpus13 (PQ963482) | Serbia narna-like virus 1<br>isolate 85061 RNA-dependent RNA polymerase<br>gene (MT822183.1) | 3280 | 2751* | 94.00 |
| XiangYun narna-levi-like virus 8<br>isolate Cx.nigripalpus14<br>(PQ963483) | XiangYun narna-levi-like<br>virus 8 isolate XY101736<br>RNA-dependent RNA<br>polymerase mRNA<br>(OL700144.1) | 2649 | 2652* | 90.61 |
| Bunyavirales sp. isolate<br>Cx.nigripalpus15 (PQ869759)<sup>†</sup> |  |  |  |  |
| Qingnian mosquito virus isolate<br>Cx.nigripalpus16 (PQ963481)<sup>†</sup> |  |  |  |  |

\* represents the length of the contig submitted for NCBI.
<sup>†</sup> represents a case where there was no best match in NCBI.

### RNA-Seq Processing Statistics: *de Novo* Assembly (Method Two)

The *de novo* assembly performed in Method Two yielded 1,187,014 contigs, with sequence lengths ranging from 172 bp to 33,372 bp. The total assembly spanned 671,847,487 bp, yielding an average contig length of 566 bp and an N50 of 726 bp. Out of total contigs 1,187,014, 695,395 contigs belong to <400 bp, 352,746 contigs ranged from 400–1000 bp, and 138,873 contigs were >1000 bp (Figure 3A). Following the Method Two pipeline, the analysis identified 56 unique contigs (∼0.004% of the total) as viral based on significant alignments; 24 of the 56 contigs had multiple BLAST hits. The number of viral contigs recovered by Method Two was ∼10-fold lower than in Method One. This difference is partly attributable to database choice: Method One performed BLAST searches against the broader NCBI nt database (2023) to maximize sensitivity, including detection of divergent or uncurated viral sequences, whereas Method Two queried the smaller, viral-only NCBI genomic database (viral.1.1.genomic.fna) to maximize specificity and reduce ambiguous matches. However, because the pipelines also differed in contig generation and filtering, differences in viral contig counts likely reflect both database coverage and contig/pipeline effects and cannot be attributed to database choice alone without cross-annotation. Among these, two contigs corresponded to viruses that were not recovered in Method One: Spodoptera litura nucleopolyhedrovirus isolate Cx.nigripalpus4 (PQ963474) and Spodoptera littoralis nucleopolyhedrovirus isolate Cx.nigripalpus5(PQ963475). The remaining 1,186,958 contigs failed to align with known viral genomes, suggesting origins from the host transcriptome, environmental contaminants, or potentially novel viral taxa. For the Merida virus, 29 individual contigs were integrated using an IUPAC consensus-based approach to account for nucleotide ambiguity across overlapping regions to assemble a viral contig to compare to Method One. Of the 56 unique contigs, 50 contigs were identified as having the keyword “virus” in their matching hits on Genbank, and six contigs had closest matches to “phage” (Figure 3C).

### Fifteen different viruses were discovered using dual methods

To find the most viral diversity in the sample, two bioinformatic methods were used and compared to try to identify the maximum number of viruses present. Cumulatively, five viruses were positive sense ssRNA viruses, three viruses were negative-sense ssRNA viruses, and eight viruses had an uncharacterized genome. Method One generated ten unique viruses like Hubei virga-like virus 3 isolate Cx.nigripalpus6 (PQ963470), Culex rhabdovirus isolate Cx.nigripalpus7 (PQ963476), Ecclesville picorna-like virus isolate Cx.nigripalpus8 (PQ963477), Elemess virus isolate Cx.nigripalpus9 (PQ963478), Elemess virus isolate Cx.nigripalpus10 (PQ963479), Qingnian mosquito virus isolate Cx.nigripalpus16 (PQ963480), Rinkaby virus isolate Cx.nigripalpus12 (PQ963485), Serbia narna-like virus 2 isolate Cx.nigripalpus13 (PQ963482), XiangYun narna-levi-like virus 8 isolate Cx.nigripalpus14 (PQ963483), Bunyavirales sp. isolate Cx.nigripalpus15 (PQ869759, name assigned by NCBI) and Qingnian mosquito virus isolate Cx.nigripalpus16 ( PQ963481, name assigned by NCBI) (Table 3).

Compared with Method Two, Method One aligned to mosquito sequences and identified viral contigs directly (Figure 3C). This method identified five total viruses with large contigs. Compared to Method One, two additional viruses were found, Spodoptera litura nucleopolyhedrovirus isolate Cx.nigripalpus5 (PQ963474) and Spodoptera littoralis nucleopolyhedrovirus isolate Cx.nigripalpus6 (PQ963475) (Table 2), indicating that each methodology has its merits. Nine contigs matched to the Spodoptera litura nucleopolyhedrovirus (NC_003102.1) and six contigs belonged to Spodoptera littoralis nucleopolyhedrovirus (NC_038369.1) within the viral BLAST dataset. These sequences were not detected in the contigs from Method One, and it is possible that these were filtered out during the mapping step in Method One, which is not present in Method Two (Table 3). Overwhelmingly, Method Two generated the most contigs matching in the NCBI viral database to either Merida virus (NC_040599.1) with twenty-nine contigs and Merida-like virus (NC_040532.1) with nine contigs. Five contigs matched to Zhejiang mosquito virus (NC_033716.1). The shortest contig found matching the viral database was 201 bp (Merida), and the longest was 9,643 bp for the Zhejiang mosquito virus. Interestingly, this virus is only 9,558 bp in GenBank, indicating that the contig identified in this study is longer than the sequence it was blasted against (Table 3). These data indicate that dual bioinformatic pipelines can add rigor, sequence coverage, and robustness to sample analyses that only utilize one methodology.

### Identification of Rhabdoviridae Members in Culex nigripalpus

Two viruses classified within the *Mononegavirales* order and *Rhabdoviridae* family were identified, both possessing non-segmented, single-stranded, negative-sense RNA genomes (Figure 4).

**Figure 4.**
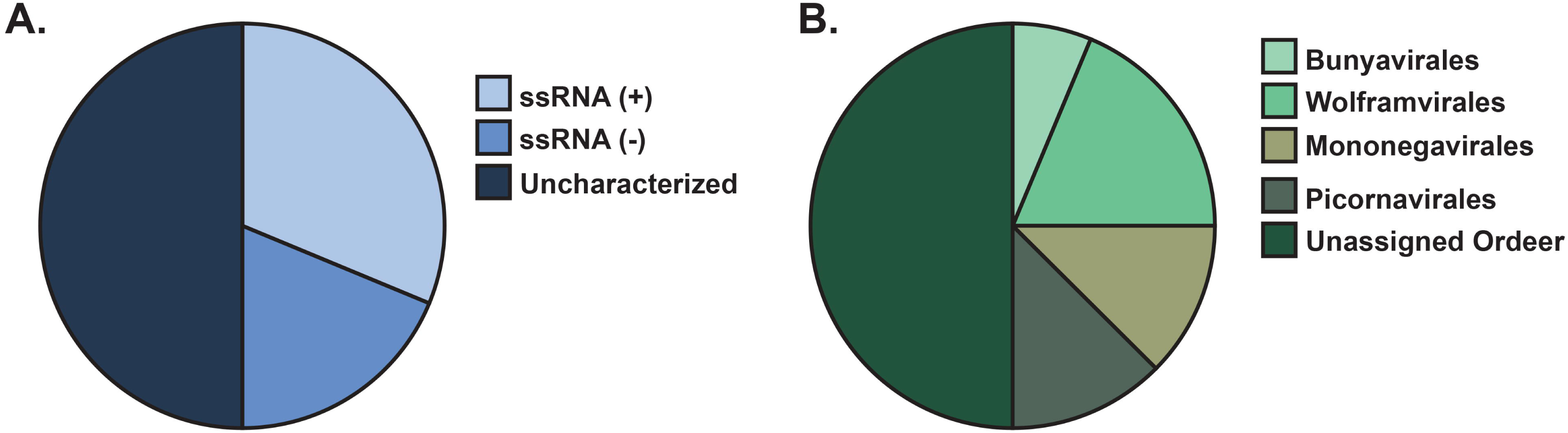
Overview of Virus diversity. A) Pie-chart representing the types of viral genomes categorized as positive-sense ssRNA, negative-sense ssRNA, or uncharacterized; ss: single-stranded; RNA: Ribonucleic acid B) Pie-chart representing virus diversity at the order level

**Figure 5.**
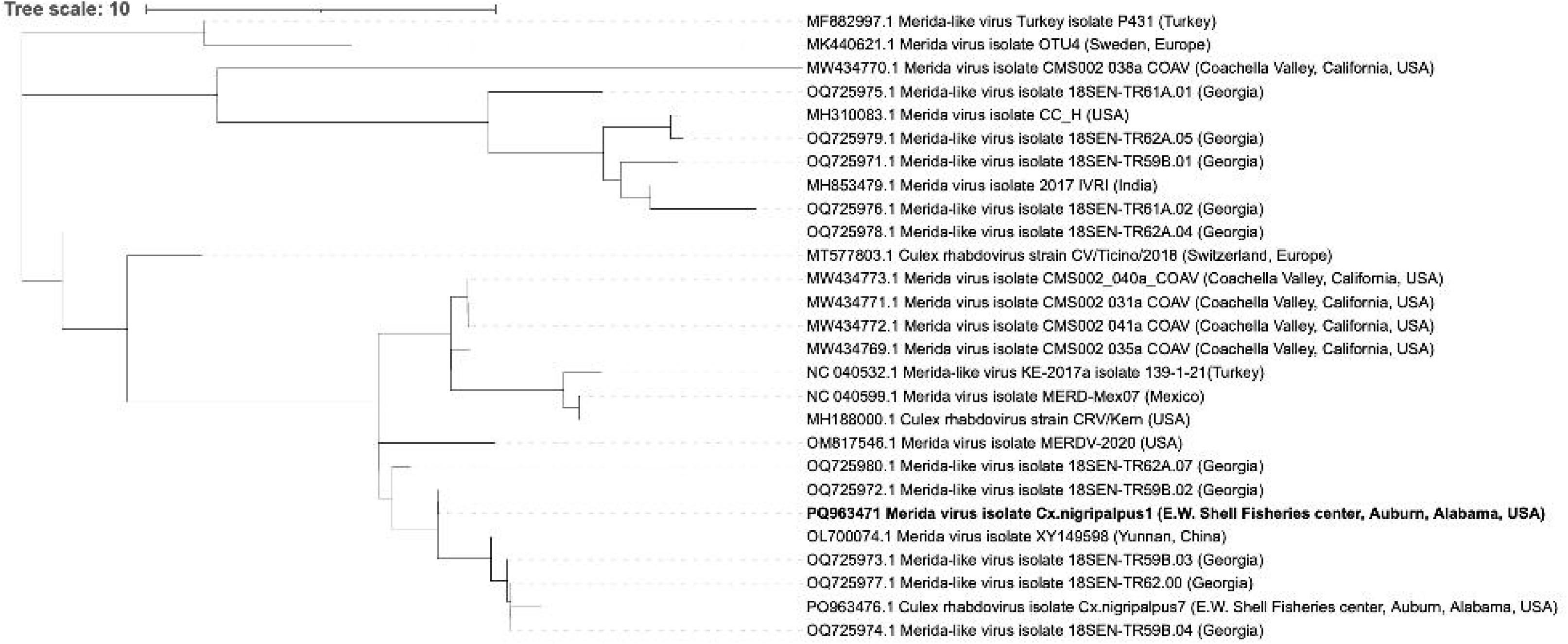
Phylogenetic analysis of Merida virus isolate Cx.nigripalpus2 (PQ963471) (Highlighted with bold lettering) assembled from Mi-Seq analysis of RNA of *Culex nigripalpus* collected in E.W. Shell Fisheries between late August to Late October in 2022

### Merida virus isolate Cx.nigripalpus1

(PQ963471) was recovered with a sequence length of 11,766 bp and exhibited 98.74% amino acid identity to Merida virus isolate MERD-Mex07 (11,798 bp, NC_040599.1). Viral coverage reached 99.72%, and five canonical genes were annotated: large protein (L), glycoprotein (G), matrix protein (M), phosphoprotein (P), and nucleoprotein (N) (Table 3, Figure 4, Supplemental Figure 1). Originally reported in *Culex quinquefasciatus* from Mexico in 2007 (Charles et al., 2016), this is the first report of a Merida virus genome in *Culex nigripalpus* and in the state of Alabama. Phylogenetic analysis (Figure 4) positioned Merida virus isolate Cx.nigripalpus2 (PQ963471) in close association with Culex rhabdovirus isolate Cx.nigripalpus7 (PQ963476), both obtained in this study. The sequence also clusters with *Merida virus MERD-Mex07* (NC_040599.1), originally detected in *Culex quinquefasciatus* from Mexico, and a U.S. isolate (MW434769.1) previously found in *Culex tarsalis* from California. This represents the first documented detection of Merida virus in *Culex nigripalpus* and in Alabama, expanding both the known host and geographic range of the virus. For Merida virus isolate Cx.nigripalpus2 (PQ963471), the normalized count for N is 220082 transcripts per million, P is 148906 transcripts per million, M is 234714 transcripts per million, G is 184730 transcripts per million, and L is 211568 transcripts per million. TPM values revealed that the M gene, despite its short length (491 bp), showed the highest normalized expression (TPM: 234,714). This suggests strong transcriptional activity during infection compared to other rhabdoviruses (Whelan et al., 2004). The RdRp gene, while having the highest estimated counts (∼1.1 million), showed moderate TPM (211,568) due to its large transcript length (6411 bp).

### Culex rhabdovirus isolate Cx.nigripalpus7 (PQ963476)

was assembled into a 3,666 bp contig, showing 98.64% amino acid identity to Culex rhabdovirus strain CRV/Kern (MH188000.1). Three complete genes were annotated: the nucleoprotein gene (116–1552 bp), phosphoprotein gene (1642–2844 bp), and matrix protein gene (2899–3459 bp), along with a partial glycoprotein segment (3568–3602 bp) (Table 3, Supplemental Figure I). This virus was previously detected in California from mosquito pools of *Culex pipiens*, *Cx. tarsalis*, and *Cx. erythrothorax* (Sadeghi et al., 2018) but has not been previously reported in Alabama.

### Partial Genome of a Serbia Narna-Like Virus Identified in *Culex nigripalpus*

A viral sequence resembling members of the order *Wolframvirales* was identified in the dataset. The contig, named Serbia narna-like virus 2 isolate Cx.nigripalpus13(PQ963482), encodes a partial RNA-dependent RNA polymerase (RdRp). It showed 94% amino acid identity to Serbian narna-like virus 1 isolate 85061 (MT822183.1) (Table 3, Supplemental Figure 1). The full genome of this virus has not yet been reported. The closest known relative is a RdRp gene of *Narnaviridae* sp. strain YX788 (MW452310.1), sharing 93.99% amino acid identity. This represents the first known detection of this virus or any close relatives in the United States. Furthermore, there are no prior reports of its presence in *Culex nigripalpus* or other mosquito species.

### Partial Picornavirales-Like Viral Genome and Unclassified Viral Fragments Identified

A partial picornavirus-like sequence related to Ecclesville picorna-like virus was identified as Ecclesville picorna-like virus isolate Cx. nigripalpus8 (PQ963477), encoding a partial RNA-dependent RNA polymerase (RdRp) (Table 3, Supplemental Figure 1). This sequence shows high amino acid similarity to strains previously isolated from *Haemagogus* mosquitoes in Trinidad (90.18%, Ali et al., 2021). This marks the first report of this virus in *Culex nigripalpus* or in the United States (Figure 4). Two additional contigs lacked strong similarity to known picornavirus genomes and were designated as distinct viral fragments: Bunyavirales sp. isolate Cx.nigripalpus15 (PQ869759) and Qingnian mosquito virus isolate Cx.nigripalpus16 (PQ963481), reflecting potentially novel or divergent viral sequences.

### Partial RdRp of Peribunyaviridae Member Detected in *Culex nigripalpus*

A partial RNA-dependent RNA polymerase (RdRp) sequence related to Qingnian mosquito virus was identified as Qingnian mosquito virus isolate Cx.nigripalpus16 (PQ963480). This contig shares 70.25% amino acid identity with Qingnian Mosquito Virus strain YC179 (MW452282.1) which was originally detected in China in 2018 (Table 2, Supplemental Figure 1). This represents the first report of a Qingnian virus-like bunyavirus in the United States and the first documented occurrence in *Culex nigripalpus*. However, based on the amino acid sequence identity, further sequencing may reveal this as a novel virus.

### Riboviria like viral sequences found in *Culex nigripalpus* in Alabama

Multiple viruses from the realm *Riboviria* were detected (Table 3, Figure 4, Supplemental Figure 1), including representatives with similarity to negeviruses, virga-like viruses, and unclassified mosquito-associated viruses from both Asia and the Americas. A partial negevirus-like genome, highly similar to Rinkaby virus originally found in Sweden, was recovered and represents the first detection of this lineage in the U.S. Similarly, a near-complete *Hubei virga-like virus 3* genome showed close identity to Californian isolates but was previously undocumented in *Culex nigripalpus*. For Hubei virga-like virus 3 isolate Cx. nigripalpus6 (PQ963470.1), the normalized transcript per million of capsid protein compared to the polyprotein is 763758, which is higher than the polyprotein (TPM: 236242). The data might suggest the indication of virus replication; however, more research is required.

Fragments of Elemess virus—including capsid and RdRp segments—were also detected, further supporting the presence of this virus beyond its original range in California. Several unclassified *Riboviria* sequences showed high similarity to viruses from China, including strains of Zhejiang mosquito virus, Hubei mosquito virus, and XiangYun narna-levi-like virus, highlighting potential global connectivity of mosquito viromes. Notably, none of these viruses have previously been reported in Alabama or in *Culex nigripalpus*, suggesting these findings may reflect either host range expansion, broader geographic distribution, or cryptic viral diversity not previously captured in public databases.

### *Lefavirales* Putative Baculovirus-Like Sequences Identified

Two sequences encoding ribonucleotide reductases were identified with similarity to *Spodoptera* nucleopolyhedroviruses, large double-stranded DNA viruses previously associated with lepidopteran hosts in China (Table 3). These viral fragments, Spodoptera litura nucleopolyhedrovirus isolate Cx.nigripalpus4 (PQ963474) and Spodoptera littoralis nucleopolyhedrovirus isolate Cx.nigripalpus5(PQ963475), shared approximately 71.7% amino acid identity to *S. litura* and *S. littoralis* nucleopolyhedroviruses, respectively (Table 2). While these viruses are known insect pathogens, their detection in *Culex nigripalpus* from Alabama is a novel finding. No prior reports of these viruses exist in mosquitoes from this region, and their low sequence similarity to known viruses may suggest that these are novel baculoviruses in mosquitoes. However, the full viral genome of either of these viruses was not recovered in our analysis suggesting that potentially only pieces of them are present.

### Comparative Recovery of Virus Genomes Across Analytical Methods

Using both methods combined, nearly complete genomes of three virus-like sequences were recovered: Merida virus (Cx.nigripalpus1), Hubei mosquito virus 5 (Cx.nigripalpus2), and Zhejiang mosquito virus (Cx.nigripalpus3). All three were identified by both analytical approaches but not at the same level of coverage (Table 3, Supplemental Figure 1). For Hubei mosquito virus 5, Method One produced a 663 bp contig with 89.58% amino acid identity to Atrato picorna-like virus 1 (MN661035.1), whereas Method Two generated a longer 9012 bp contig with 77.16% identity to Zhejiang mosquito virus 1 (NC033716.1). Through cross-BLAST analysis, the shorter contig was found to overlap with the longer contig; therefore, the longer sequence was selected for NCBI submission, and the name of the virus that matched the longer contig was selected.

In addition to overlapping viruses identified by both pipelines, the *de novo* assembly approach (Method Two) recovered 15 distinct viral taxa. Notably, it uncovered viral sequences that were absent from the reference-based pipeline. These included two lepidopteran baculoviruses—*Spodoptera litura* nucleopolyhedrovirus, and *Spodoptera littoralis* nucleopolyhedrovirus—as well as Merida-like virus KE-2017a, which shares high identity with

Merida virus but constitutes a distinct strain. Bacteriophage sequences were also recovered, including Escherichia phage 500465-1, Klebsiella phage ST15-OXA48phi14.1, and Klebsiella phage ST437-OXA245phi4.1. Additional hits included *Bracoviriform indiense*, likely representing a bracovirus element, Mosquito X virus, and Prymnesium kappa virus, an algal virus likely acquired from the environment. In another instance of Hubei mosquito virus 5, Method One yielded a 265 bp contig aligning from position 81 bp to 345 bp with Hubei mosquito virus 4 (4971 bp, NC_032231.1), while Method Two produced a 3734 bp contig aligning from position 1 to 3643 bp with 89.45% amino acid identity to the same virus. The 3734 bp sequence was chosen for submission based on contig length and alignment. Method Two consistently generated longer contigs of some viruses that encompassed the sequences identified by Method One, providing more complete genome assemblies. Although Method One produced slightly higher amino acid identity in one case, the extended coverage offered by Method Two enabled the selection of its contigs for NCBI submission when the sequences were directly compared. These findings indicate that Method Two had advantages as it generated longer contigs of specific viruses; however, it did not identify all the viruses found in Method One, and Method One had the advantage of identifying more unique viral sequences than Method Two (Table 3).

### Characterization of Additional Pathogen Sequences within the Dataset

Using a Plasmodium reference panel, no identified similarity to any of the Method Two *Cx. nigripalpus* contigs were found, indicating that plasmodia were not present in these insects. As no incidences of *Plasmodium hermani* (Santiago-Alarcon et al., 2012) were reported locally, this result was not surprising. To further identify sequences from Method One and Two, the unannotated sequences from the BLASTN analysis were further scrutinized using taxonomic methods to identify those with a viral designation using MEGAN. Once identified, these sequences were used in a separate BLASTX run to obtain more information. From Method One, 78 additional sequences were identified. This had 430 rows of hits in the BLASTX. Matching these against the fasta sequences, 73 sequences had hits, 66 had duplicates. Of the 73 that had hits, once sorted by the longest alignment length, highest bit score, and lowest e-value, 58 were still retained as “viral”; 15 had top hits that were insect or random sequences. (Supplemental Table II). In Method Two, there were 166 additional sequences sent in for a total of 717 BLASTX hits. Of these, 151 queries were unique. Interestingly, 14 sequences had no hits. Once sorted, 123 were retained as viral hits, and 28 were sorted out as insect or random sequences (Supplemental Table III). Interestingly, while some novel viruses were detected, the dataset was dominated by entries for Hubei virga-like virus, Merida virus, and Zhejiang mosquito virus, all of which had already been identified using BLASTN. This suggests that viral diversity within these sequences prevented complete filtering in earlier steps but did not ultimately expand the dataset.

### Validation of Viruses

After RT-PCR amplification, eleven out of twelve contigs selected for validation produced bands consistent with the sizes predicted by PrimerBlast. For Rinkaby virus, one contig with an expected amplicon size of 1109 nt produced a noticeably larger band, approximately 1500 nt. This isolated discrepancy suggests that the assembled contig was missing a region that the RT-PCR successfully recovered. All other Rinkaby virus contigs amplified at their predicted lengths, indicating the issue was limited to that single contig. Merida virus showed the expected ∼1974 nt product, aligning well with the ∼2 kb band on the gel ladder. XiangYun virus also matched predictions, with all three primer sets generating products of the correct sizes. Expected bands were similarly obtained for Elemess, Ecclesville, Hubei, and the Qingnian virus–like bunyavirus contig (Figure 6). Amplification differed between reverse transcriptase enzymes for only one target: the first Elemess virus contig amplified with RevertAid but not with the Verso cDNA mix. The most likely explanation is that differences in buffer composition, priming efficiency, or enzyme performance between the two reverse transcriptase systems affected cDNA yield for this specific target. The second Elemess contig amplified with both enzymes, indicating that this effect was limited to a single amplicon.

**Figure 6.**
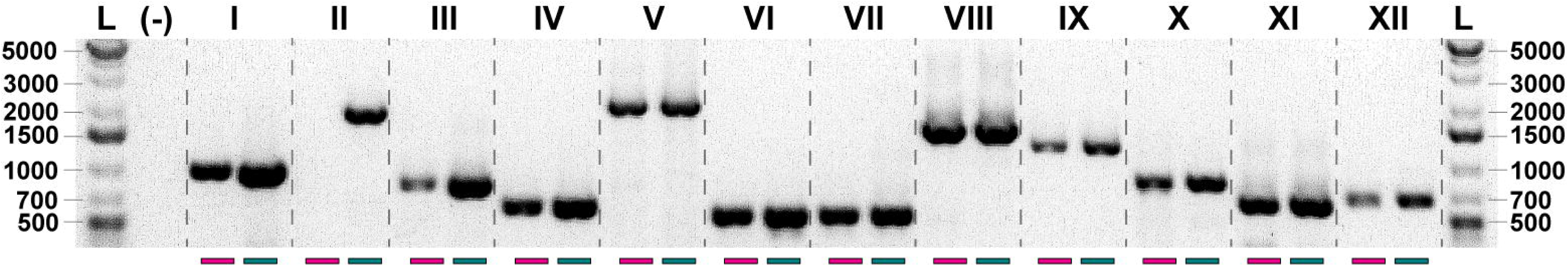
Validation of contigs from the mosquito virome analysis. Agarose gel electrophoresis of RT–PCR amplicons confirming selected viral contigs from *Culex nigripalpus*. Lanes are as follows: L, O’GeneRuler 1 kb Plus DNA ladder (Thermo Fisher Scientific); –, negative control; I, Hubei virus; II, Elemess virus; III, XiangYun virus; IV, XiangYun virus; V, Merida virus; VI, Elemess virus; VII, Ecclesville virus; VIII, Rinkaby virus; IX, XiangYun virus; X, Bunyavirales contig; XI, Rinkaby virus; XII, Rinkaby virus. Lane numbers (I–XII) correspond to primer sets listed in Supplemental Table 1. Pink bars beneath the gel indicate RT–PCRs performed with Verso, and teal bars indicate RT–PCRs performed with RevertAid.

## Discussion

This study identified *Culex nigripalpus* as the predominant mosquito species amplified at an Alabama aquaculture facility and characterized its associated virome. To interrogate viral content in the RNA-Seq dataset, we applied two complementary bioinformatics pipelines that differed in how host sequences were handled and in how contigs were assembled and annotated. In the reference-based assembly pipeline (Method One), reads were mapped to the *Culex quinquefasciatus* genome and only unmapped reads were assembled *de novo* and queried against the NCBI nt database. This strategy enriches for non-host sequences but can discard viral reads that share homology with the reference genome or occur at low abundance, particularly when a closely related rather than conspecific genome is used (Ekblom and Wolf, 2014; MacManes, 2016). In the second pipeline using *de novo* assembly (Method Two), all reads were assembled *de novo* and contigs were queried directly against the NCBI viral genome database. *De novo* assembly can facilitate the detection of divergent or novel viruses by reconstructing longer contigs that align even when no close reference genome is available, as shown in large-scale invertebrate virome surveys (Shi et al., 2016). Still, it also increases the risk of chimeric contigs and other assembly artifacts that can result from contaminant host reads (Vollmers et al., 2017) and remains constrained by incomplete viral reference databases. Databases that underrepresent many environmental and arthropod-associated viruses (Rosario and Breitbart, 2011; Simmonds et al., 2017), leaving some true viral sequences unclassified or only assignable at higher taxonomic levels. The overlap and differences between these pipelines underscore their complementary strengths, and we recommend that future metatranscriptomic studies adopt multiple analytical approaches when possible to maximize information gained from labor-intensive field collections.

Here we defined the mosquito virome broadly as any sequences of viral origin detected in the pooled *Cx. nigripalpus* sample, including actively replicating viruses, viruses associated with the mosquito microbiome, and viral fragments that may be integrated into the mosquito genome (Bonning, 2019). Consistent with the ecology of the site and the known host range of *Cx. nigripalpus*, nearly all viruses detected were insect-specific or not known to infect humans. The absence of recognized human-pathogenic arboviruses aligns with this species’s primarily ornithophilic feeding behavior, although it will opportunistically bite mammals, including humans, when available. These results illustrate how mosquito behavior and habitat structure the virome and show that fisheries-adjacent to semi-natural environments can harbor viral diversity that is not captured in urban, human-focused surveillance programs. The viruses identified in *Cx. nigripalpus* span a broad taxonomic range, with many RNA viruses drawn from multiple orders. Such diversity is consistent with previous mosquito virome surveys and suggests a stable association between *Cx. nigripalpus* and a diverse assemblage of insect-specific viruses, providing a useful baseline for future comparative and longitudinal studies. Insect-specific viruses have been implicated in modulating mosquito susceptibility to medically important arboviruses, so defining this background virome in *Cx. nigripalpus* provides a foundation for future work on how these persistent infections may influence vector competence in aquaculture-linked habitats.

Both pipelines depend on sequence similarity searches against public databases, which represent a major bottleneck for virome studies. Insect-associated viruses remain underrepresented in NCBI nt and viral genome collections, so sequences from highly divergent or truly novel viruses may fail to be annotated and were likely present among the many unclassified contigs in our assemblies. Our additional DIAMOND–MEGAN analysis recovered relatively few extra viral taxa. It was dominated by the same Hubei virga-like, Merida, and Zhejiang mosquito virus sequences identified by BLASTN, highlighting this limitation. Proteomics and other orthogonal approaches could help validate viral expression independent of sequence similarity, although peptide identification ultimately still relies on underlying sequence data (Baldridge et al., 2016). Continued development of homology-independent algorithms, long-read sequencing, and targeted virus-enrichment strategies will be important for reducing these detection biases; the present study provides a foundation for such work in *Cx. nigripalpus*.

Among the RNA viruses detected, we focused in particular on RNA-dependent RNA polymerase (RdRp) sequences, which are central to viral replication and transcription (Venkataraman et al., 2018). Five distinct RdRp-encoding viruses were identified, and three of these—Serbia narna-like virus 2, XiangYun narna-levi-like virus 8, and Elemess virus—retained conserved catalytic motifs previously shown to be essential for polymerase function (Jia and Gong, 2019). Recovery of these conserved domains may reflect higher transcript abundance, greater stability, or improved detectability of these sequences during assembly, implying that viruses with strongly conserved polymerase motifs are more likely to be captured in RNA-Seq datasets. Conversely, viruses with highly divergent polymerases or alternative replication strategies may be systematically underrepresented.

Many of the viruses identified here have been reported previously from other mosquito species or locations, but not from *Cx. nigripalpus* in Alabama. Merida virus, Hubei virga-like virus 3, and Elemess virus have all been detected in North America in other Culex species; whereas several additional viruses showed closest matches to sequences reported from China (Shi et al., 2016), Serbia (Stanojević et al., 2020), Sweden, and the Mediterranean region. Ecclesville picorna-like virus, for example, was originally described from Haemagogus mosquitoes in a bird sanctuary in Trinidad and Tobago (Ali et al., 2021). Members of the Picornavirales have broad host ranges (Zell et al., 2017), so this virus may be associated with avian or aquatic hosts at our site and retained in *Cx. nigripalpus* after blood feeding. The E.W. Shell Fisheries Center combines aquaculture ponds with surrounding forest and open habitats, exposing *Cx. nigripalpus* to birds, reptiles, amphibians, and other vertebrates, as well as occasional human hosts. This diverse host community likely contributes to the breadth of viruses detected.

Collectively, these observations support the idea that mosquito-associated viruses can be geographically widespread, with closely related lineages occurring in distinct ecological settings and host assemblages. Merida virus has previously been reported from *Cx. tarsalis* and *Cx. quinquefasciatus* in the United States, yet the Merida virus isolate Cx. nigripalpus2 (PQ963471) recovered here is most similar to the strain first identified in *Cx. quinquefasciatus* from Mexico in 2007 (Charles et al., 2016). This pattern is consistent with long-distance dispersal, potentially mediated by weather events that can reshape insect communities and vector movement (Novais et al., 2018). Regardless of its precise origin, our data document Merida virus in *Cx. nigripalpus* and in Alabama for the first time, expanding both the host and geographic range of this virus. The near-complete genome assemblies recovered for Merida virus (11,766 of 11,798 bp) and Hubei virga-like virus 3 (10,813 of 10,870 bp), together with high transcript abundances, suggest that these viruses were among the most transcriptionally active or abundant in the sampled population.

The dual-pipeline strategy also improved detection of viruses with different genomic properties. Three viruses: Merida virus, Hubei mosquito virus 5, and Zhejiang mosquito virus 1, were recovered by both pipelines, providing reciprocal support for their presence and yielding more complete genome reconstructions. In contrast, the *de novo*–only pipeline uniquely detected two baculovirus-like sequences related to Spodoptera litura nucleopolyhedrovirus and Spodoptera littoralis nucleopolyhedrovirus (Agboli et al., 2019; Williams et al., 2017). These fragments encoded ribonucleotide reductases and may represent endogenous viral elements (EVEs), which are well documented for retroviruses (Hindmarsh and Leis, 1999) and increasingly recognized among non-retroviral taxa, including flaviviruses and rhabdoviruses (Crochu et al., 2004; Katzourakis and Gifford, 2010; Whitfield et al., 2017). Whether these baculovirus-like sequences reflect integration events, environmental contamination, or transient acquisition from lepidopteran prey or larvae remains unresolved and merits further study.

Overall, these results emphasize that virome composition is shaped by local ecology and that interpretation of viral detection data must account for the behavior and habitat of the vector. Although we identified multiple viruses associated with *Cx. nigripalpus* at a fisheries-adjacent site, their ecological and evolutionary roles remain unresolved. Future work should address whether these viruses are vertically transmitted, persist in environmental reservoirs, or interact with one another in ways that alter mosquito fitness or vector competence. This study was based on pooled mosquitoes from a single location and time period, so broader spatial and temporal sampling, including individual-level analyses, will be important for understanding how generalizable these patterns are across Alabama and beyond. Despite these limitations, the complementary bioinformatic approaches used here provided a more complete picture of the *Cx. nigripalpus* virome than either would alone and offer a framework for future studies of virus–vector–environment interactions.

## Conclusion

This study provides the first characterization of the *Culex nigripalpus* virome in Alabama and shows that aquaculture-associated habitats can amplify this species and support a diverse assemblage of primarily insect-specific viruses. Using RNA-Seq, we detected at least fifteen distinct viruses associated with *Cx. nigripalpus* at a fisheries-adjacent site, including multiple first reports for this host and region. Most notably, we identified a Merida virus isolate (GenBank accession PQ963471), representing the first documentation of this virus in both Alabama and in association with *Cx. nigripalpus*, along with several viruses previously known only from other continents or mosquito hosts. These findings highlight *Cx. nigripalpus* as a relevant target for arboviral and insect-specific virus surveillance in aquaculture-linked landscapes. Methodologically, the comparison of a reference-based pipeline and a de novo assembly–based pipeline showed that each approach recovers overlapping but non-identical sets of viruses; the dual strategy improved genome completeness for several taxa and revealed additional viral and viral-like sequences, including putative baculovirus fragments that may represent endogenous viral elements or environmentally derived DNA. Together, these results underscore the importance of integrating complementary bioinformatic workflows to maximize viral discovery and genome recovery from metatranscriptomic datasets and provide a framework for future work examining how aquaculture, local ecology, and mosquito biology interact to shape virome composition and vector–virus dynamics in the southeastern United States.

## Supporting information

Supplemental Table 1_Primers

Supplemental Table III

## Supplemental

Supplemental Figure 1: Virus Map: Graphical representation viruses from Mi-Seq of Mosquito (*Culex nigripalpus*) collected in Auburn, Alabama in 2022; Viruses belonging to (A) Mononegavirales (B) Wolframvirales (C) Picornavirales (D) Bunyavirales (E) Patatavirales (F) Riboviria realm (G) Lefavirales (H) Hypothetical viruses; ORF: Open reading frames; RdRP: RNA-dependent RNA polymerase; NCBI: National Center for Biotechnology Information. The scale is reflective of the amount of sequence from this project and not necessarily reflective of the whole genome of the virus in question. When possible, reference sequences were used to anchor this study’s sequences to a reference to gain perspective about how much was sequenced compared to what was expected.

Supplemental Table I: Primer sequences used for validation of contigs of virome from *Culex nigripalpus*.

Supplemental Table 1I: Best hits from DIAMOND/MEGAN from Method One.

Supplemental Table III: Best hits from DIAMOND/MEGAN from Method Two.

Supplemental File 1: Fasta file of COI sequences from *Cx. nigripalpus*.

Supplemental File 2: Fasta file of sequences of Method One retrieved after DIAMOND/MEGAN which were used for BLASTX shown in Supplmental Table I.

Supplemental File 3: Fasta file of sequences of Method Two retrieved after DIAMOND/MEGAN which were used for BLASTX shown in Supplmental Table II.

## References

Agboli, E., Leggewie, M., Altinli, M., Schnettler, E., 2019. Mosquito-Specific Viruses—Transmission and Interaction. Viruses 11, 873.

Ali, R., Jayaraj, J., Mohammed, A., Chinnaraja, C., Carrington, C.V.F., Severson, D.W., Ramsubhag, A., 2021. Characterization of the virome associated with Haemagogus mosquitoes in Trinidad, West Indies. Scientific Reports 11, 16584.

Bailey, M.A., 1998. Snakes of Alabama: Fact and fiction. Alabama’s Treasured Forests, 28–30.

Baldridge, G.D., Markowski, T.W., Witthuhn, B.A., Higgins, L., Baldridge, A.S., Fallon, A.M., 2016. The Wolbachia WO bacteriophage proteome in the Aedes albopictus C/w Str1 cell line: evidence for lytic activity? In Vitro Cellular & Developmental Biology-Animal 52, 77–88.

Beckmann, J.F., Fallon, A.M., 2012. Decapitation improves detection of Wolbachia pipientis (Rickettsiales: Anaplasmataceae) in Culex pipiens (Diptera: Culicidae) mosquitoes by the polymerase chain reaction. J Med Entomol 49, 1103–1108.

Bonning, B.C., 2019. The Insect Virome: Opportunities and Challenges, Current Issues in Molecular Biology, pp. 1–12.

Bray, N.L., Pimentel, H., Melsted, P., Pachter, L., 2016. Near-optimal probabilistic RNA-seq quantification. Nature Biotechnology 34, 525–527.

Burkett-Cadena, N.D., Blosser, E.M., Reeves, L.E., 2022. Key to the Adult Females of Species of Culex Subgenus Melanoconion in Florida, USA. Journal of the American Mosquito Control Association 38, 130–140.

Burr, P.C., Avery, J.L., Street, G.M., Strickland, B.K., Dorr, B.S., 2020. Historic and contemporary use of catfish aquaculture by piscivorous birds in the Mississippi Delta. The Condor: Ornithological Applications 122, duaa036.

Charles, J., Firth, A.E., Loroño-Pino, M.A., Garcia-Rejon, J.E., Farfan-Ale, J.A., Lipkin, W.I., Blitvich, B.J., Briese, T., 2016. Merida virus, a putative novel rhabdovirus discovered in Culex and Ochlerotatus spp. Mosquitoes in the Yucatan Peninsula of Mexico. Journal of General Virology 97, 977–987.

Community, T.G., 2024. The Galaxy platform for accessible, reproducible, and collaborative data analyses: 2024 update. Nucleic Acids Research 52, W83–W94.

Crochu, S., Cook, S., Attoui, H., Charrel, R.N., De Chesse, R., Belhouchet, M., Lemasson, J.-J., de Micco, P., de Lamballerie, X., 2004. Sequences of flavivirus-related RNA viruses persist in DNA form integrated in the genome of Aedes spp. mosquitoes. Journal of General Virology 85, 1971–1980.

Darsie, R.J., Ward, R.A., 2005. Identification and Geographical Distribution of the Mosquitoes of North America, North of Mexico. University Press of Florida, Gainesville, FL.

Deng, X., Naccache, S.N., Ng, T., Federman, S., Li, L., Chiu, C.Y., Delwart, E.L., 2015. An ensemble strategy that significantly improves de novo assembly of microbial genomes from metagenomic next-generation sequencing data. Nucleic Acids Research 43, e46–e46.

Deng, Z., Delwart, E., 2021. ContigExtender: a new approach to improving de novo sequence assembly for viral metagenomics data. BMC Bioinformatics 22, 119.

Ekblom, R., Wolf, J.B.W., 2014. A field guide to whole-genome sequencing, assembly and annotation. Evolutionary Applications 7, 1026–1042.

Forrester, D.J., Nayar, J.K., Foster, G.W., 1980. Culex nigripalpus: a natural vector of wild turkey malaria (Plasmodium hermani) in Florida. J Wildl Dis 16, 391–394.

Godsey, M.S., Jr., King, R.J., Burkhalter, K., Delorey, M., Colton, L., Charnetzky, D., Sutherland, G., Ezenwa, V.O., Wilson, L.A., Coffey, M., Milheim, L.E., Taylor, V.G., Palmisano, C., Wesson, D.M., Guptill, S.C., 2013. Ecology of potential West Nile virus vectors in Southeastern Louisiana: enzootic transmission in the relative absence of Culex quinquefasciatus. Am J Trop Med Hyg 88, 986–996.

Grabherr, M.G., Haas, B.J., Yassour, M., Levin, J.Z., Thompson, D.A., Amit, I., Adiconis, X., Fan, L., Raychowdhury, R., Zeng, Q., Chen, Z., Mauceli, E., Hacohen, N., Gnirke, A., Rhind, N., di Palma, F., Birren, B.W., Nusbaum, C., Lindblad-Toh, K., Friedman, N., Regev, A., 2011. Full-length transcriptome assembly from RNA-Seq data without a reference genome. Nature Biotechnology 29, 644–652.

Hassan, H.K., Cupp, E.W., Hill, G.E., Katholi, C.R., Klingler, K., Unnasch, T.R., 2003. Avian host preference by vectors of eastern equine encephalomyelitis virus. Am J Trop Med Hyg 69, 641–647.

Hindmarsh, P., Leis, J., 1999. Retroviral DNA Integration. Microbiology and Molecular Biology Reviews 63, 836–843.

Jia, H., Gong, P., 2019. A Structure-Function Diversity Survey of the RNA-Dependent RNA Polymerases From the Positive-Strand RNA Viruses. Frontiers in Microbiology Volume 10–2019.

Katzourakis, A., Gifford, R.J., 2010. Endogenous Viral Elements in Animal Genomes. PLOS Genetics 6, e1001191.

Koh, C., Frangeul, L., Blanc, H., Ngoagouni, C., Boyer, S., Dussart, P., Grau, N., Girod, R., Duchemin, J.-B., Saleh, M.-C., 2023. Ribosomal RNA (rRNA) sequences from 33 globally distributed mosquito species for improved metagenomics and species identification. eLife 12, e82762.

Kumar, S., Stecher, G., Suleski, M., Sanderford, M., Sharma, S., Tamura, K., 2024. MEGA12: Molecular Evolutionary Genetic Analysis Version 12 for Adaptive and Green Computing. Molecular Biology and Evolution 41.

MacManes, M.D., 2016. Establishing evidenced-based best practice for the *de novo* assembly and evaluation of transcriptomes from non-model organisms. bioRxiv, 035642.

Mount, R.H., 1975. The Reptiles and Amphibians of Alabama. Auburn University Agricultural Experiment Station, Auburn, Alabama.

Novais, S., Macedo-Reis, L.E., Cristobal-Peréz, E.J., Sánchez-Montoya, G., Janda, M., Neves, F., Quesada, M., 2018. Positive effects of the catastrophic Hurricane Patricia on insect communities. Scientific Reports 8, 15042.

Payá-Milans, M., Olmstead, J.W., Nunez, G., Rinehart, T.A., Staton, M., 2018. Comprehensive evaluation of RNA-seq analysis pipelines in diploid and polyploid species. GigaScience 7.

Qualls, W.A., Mullen, G.R., 2006. Larval survey of tire-breeding mosquitoes in Alabama. J Am Mosq Control Assoc 22, 601–608.

Richards, S.L., Anderson, S.L., Lord, C.C., Tabachnick, W.J., 2011. Impact of West Nile virus dose and incubation period on vector competence of Culex nigripalpus (Diptera: Culicidae). Vector Borne Zoonotic Dis 11, 1487–1491.

Rosario, K., Breitbart, M., 2011. Exploring the viral world through metagenomics. Current Opinion in Virology 1, 289–297.

Santiago-Alarcon, D., Palinauskas, V., Schaefer, H.M., 2012. Diptera vectors of avian Haemosporidian parasites: untangling parasite life cycles and their taxonomy. Biological Reviews 87, 928–964.

Shaikh, A.A., Johnson, W.E., Jr., Stevens, C., Tang, A.Y., 1987. The isolation of spiroplasmas from mosquitoes in Macon County, Alabama. J Am Mosq Control Assoc 3, 289–295.

Shi, M., Lin, X.-D., Tian, J.-H., Chen, L.-J., Chen, X., Li, C.-X., Qin, X.-C., Li, J., Cao, J.-P., Eden, J.-S., Buchmann, J., Wang, W., Xu, J., Holmes, E.C., Zhang, Y.-Z., 2016. Redefining the invertebrate RNA virosphere. Nature 540, 539–543.

Simmonds, P., Adams, M.J., Benkő, M., Breitbart, M., Brister, J.R., Carstens, E.B., Davison, A.J., Delwart, E., Gorbalenya, A.E., Harrach, B., Hull, R., King, A.M.Q., Koonin, E.V., Krupovic, M., Kuhn, J.H., Lefkowitz, E.J., Nibert, M.L., Orton, R., Roossinck, M.J., Sabanadzovic, S., Sullivan, M.B., Suttle, C.A., Tesh, R.B., van der Vlugt, R.A., Varsani, A., Zerbini, F.M., 2017. Virus taxonomy in the age of metagenomics. Nature Reviews Microbiology 15, 161–168.

Stanojević, M., Li, K., Stamenković, G., Ilić, B., Paunović, M., Pešić, B., Maslovara, I.Đ., Šiljić, M., Ćirković, V., Zhang, Y., 2020. Depicting the RNA Virome of Hematophagous Arthropods from Belgrade, Serbia, Viruses.

Stecher, G., Suleski, M., Tao, Q., Tamura, K., Kumar, S., 2025. MEGA 12.1: Cross-Platform Release for macOS and Linux Operating Systems. J Mol Evol.

Venkataraman, S., Prasad, B.V.L.S., Selvarajan, R., 2018. RNA Dependent RNA Polymerases: Insights from Structure, Function and Evolution. Viruses 10, 76.

Vollmers, J., Wiegand, S., Kaster, A.-K., 2017. Comparing and Evaluating Metagenome Assembly Tools from a Microbiologist’s Perspective - Not Only Size Matters! PLOS ONE 12, e0169662.

Whelan, S.P., Barr, J.N., Wertz, G.W., 2004. Transcription and replication of nonsegmented negative-strand RNA viruses. Curr Top Microbiol Immunol 283, 61–119.

Whitfield, Z.J., Dolan, P.T., Kunitomi, M., Tassetto, M., Seetin, M.G., Oh, S., Heiner, C., Paxinos, E., Andino, R., 2017. The Diversity, Structure, and Function of Heritable Adaptive Immunity Sequences in the Aedes aegypti Genome. Current Biology 27, 3511–3519.e3517.

Williams, T., Virto, C., Murillo, R., Caballero, P., 2017. Covert Infection of Insects by Baculoviruses. Frontiers in Microbiology Volume 8–2017.

Zell, R., Delwart, E., Gorbalenya, A.E., Hovi, T., King, A.M.Q., Knowles, N.J., Lindberg, A.M., Pallansch, M.A., Palmenberg, A.C., Reuter, G., Simmonds, P., Skern, T., Stanway, G., Yamashita, T., Consortium, I.R., 2017. ICTV Virus Taxonomy Profile: Picornaviridae. Journal of General Virology 98, 2421–2422.

Zhao, C., Escalante, C., Jacobson, A.L., Balkcom, K.S., Conner, K.N., Martin, K.M., 2025. Metatranscriptomic and metagenomic analyses of cotton aphids (Aphis gossypii) collected from cotton fields in Alabama, USA. Frontiers in Insect Science Volume 5 - 2025.

