## Supplementary figures and images for "Virome of *Culex nigripalpus* at an Alabama aquaculture site reveals diverse insect-specific viruses and the impact of dual bioinformatic pipelines"

### Supplemental Figure 1

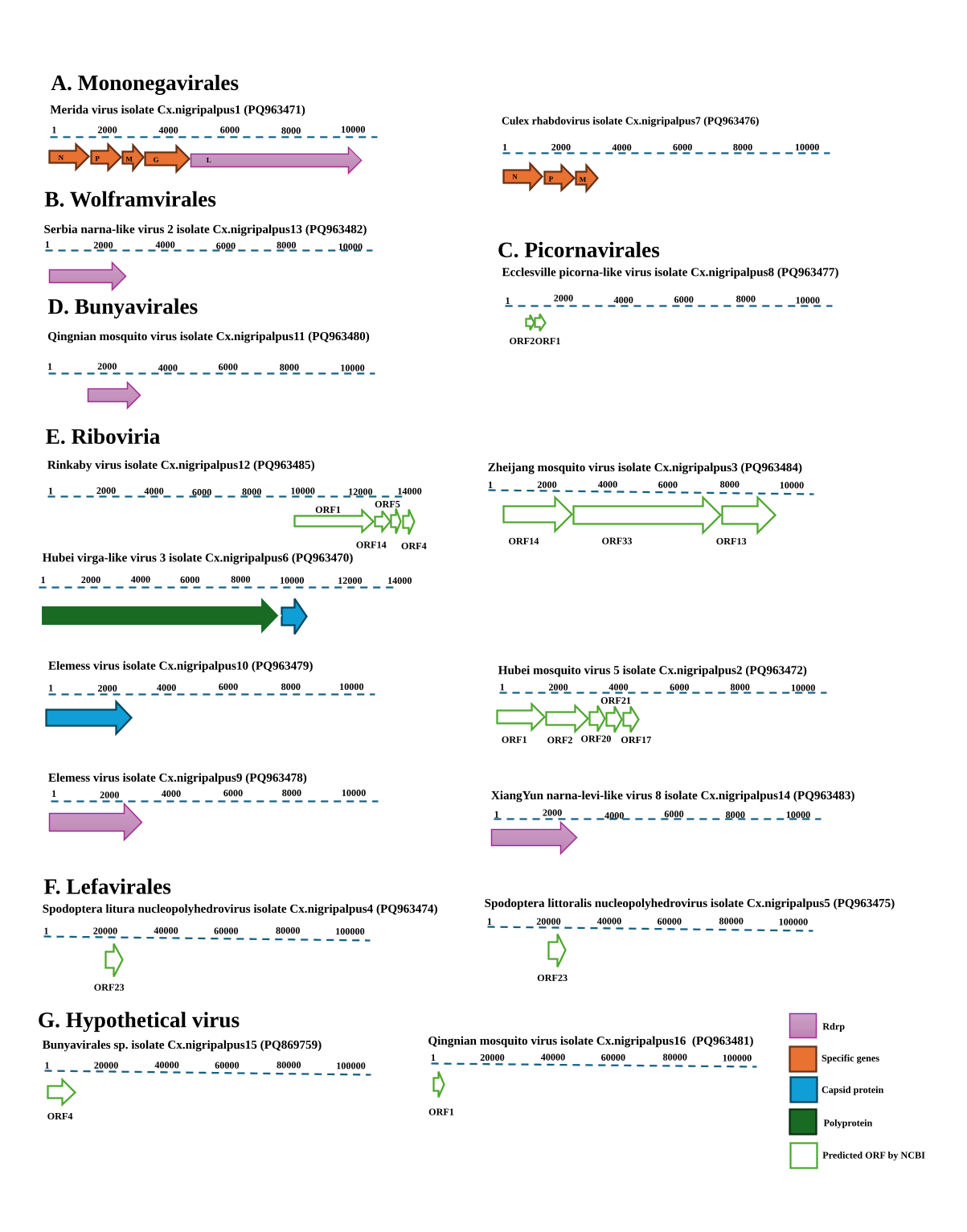
